# Methane oxidation genes are only present in bioaerosols at sites with high methane concentrations

**DOI:** 10.64898/2026.10.13.758901

**Authors:** Jungsoo Park, Nesil Han Martens, Sascha M.B. Krause, Anand Kumar, Naomi Zimmerman, Nadine Borduas-Dedekind, S. Jane Fowler

**Affiliations:** Department of Biological Sciences, Simon Fraser University, Burnaby, BC, Canada; National Field Observation and Research Station (Zhejiang Tiantong) for Forest Ecosystem, School of Ecological and Environmental Sciences, East China Normal University, Shanghai, China; Department of Mechanical Engineering, University of British Columbia, Vancouver, BC, Canada; Department of Chemistry, University of British Columbia, Vancouver, BC, Canada

## Abstract

Airborne microorganisms connect Earth’s habitats, yet a central question in microbial ecology remains: do airborne communities simply reflect what is released from the surfaces below, or does local environmental context shape them? Methane oxidation offers a tractable test, because methane monooxygenase genes are restricted to defined bacterial lineages and methane is directly measurable. Pairing aerosol and underlying soil or water metagenomes across a 35-fold methane range, we show that these genes were present in every soil and water sample, yet detected in aerosols only above the landfill and dairy farm, and not above the three wetlands. The soil did not predict the air; instead, gene detection tracked local methane availability. Aerosol cell concentrations spanned 10³ to 10⁶ cells m⁻³, and bacterial communities tracked their sources whereas eukaryotes were enriched in spore-dispersing fungi. Predicting what airborne microbiomes carry will therefore require accounting for local environmental context, not passive dispersal alone.

## Introduction

Microorganisms are ubiquitous in atmospheric aerosols, yet a central open question in microbial ecology is whether the atmosphere is a passive corridor, in which airborne communities simply reflect what is released from surfaces, or whether air acts as a selective environment, shaping communities during aerosolization (the release of cells from surfaces into air) and subsequent airborne residence ^1–3^. Aerosolized microbes are exposed to environmental stressors, including ultraviolet radiation, an oxidizing environment, desiccation, and nutrient limitation ^4,5^. Airborne microbial cell concentrations are substantially lower than those reported for soils and aquatic systems ^6^, and many detected taxa can be traced to adjacent terrestrial and aquatic microbiomes ^3^. The long prevailing view among environmental microbiologists is the first: air is generally not considered a habitat, and airborne microbes are assumed to be a passive sample of their sources. Under this passive dispersal model, each taxon, and the genes it carries, enters the air in proportion to its abundance at the source. This assumption has rarely been tested directly, because testing it requires aerosol metagenomes paired with their own source communities together with a measured variable that predicts which functions should differ. This study provides such a test, examining whether local environmental conditions, in this case the availability of a growth substrate, shape the taxa and functional genes found in near-source aerosols, sampled directly above their source environments.

Several lines of evidence indicate that atmospheric residence imposes selective pressures beyond proportional dispersal from source communities ^7^. Airborne bacterial and fungal communities in isolated Antarctic habitats show limited compositional overlap with local soils, indicating that airborne and soil communities remain distinct even within a region ^8^. Similarly, recurrent seasonal patterns in airborne bacterial communities indicate that certain taxa are consistently detected over time, consistent with structured recurrence rather than stochastic dispersal alone ^9^. Some taxa possess traits that confer resistance to ultraviolet radiation, oxidative stress, and desiccation ^10–13^, and such traits may bias community composition during airborne residence. Beyond stress tolerance, survival under the energy-limited conditions of the atmosphere can be supported by oxidation of trace gases including hydrogen, carbon monoxide, and methane at near-atmospheric concentrations, sustaining maintenance metabolism during organic carbon deprivation ^14–17^. These findings suggest that the ability to exploit diverse substrates for energy metabolism may confer a selective advantage during atmospheric residence, although this has not yet been demonstrated.

Characterizing functional gene representation in aerosols has remained technically challenging because of the extremely low microbial biomass present in air samples. Collecting sufficient biomass requires sampling large air volumes, and DNA yields are frequently near detection limits, increasing potential contributions of contamination and sequencing noise ^18,19^. Early shotgun metagenomic studies of airborne microbiomes relied on whole-genome amplification to compensate for low DNA yields, introducing biases that complicate community-level interpretation ^19^. As a result, most aeromicrobiome research has relied on 16S rRNA gene amplicon surveys, which provide robust taxonomic profiles but limited resolution of metabolic gene repertoires ^3,20^. Although optimized protocols for low-biomass aerosol sampling and DNA extraction are now available ^21^, environmentally paired shotgun metagenomic datasets that include both aerosol and source community profiles remain uncommon. To address this, we apply a gene-centric framework using curated marker genes defined *a priori* from canonical metabolic pathways, providing a conservative and reproducible strategy for comparing functional repertoires across paired aerosol and environmental metagenomes.

Methane oxidation by methane-oxidizing bacteria (methanotrophs) provides a tractable model for testing context-dependent gene retention in aerosols. The physiology, ecology and genomics of methanotrophs are well characterized ^22–24^, and they are key players in regulating methane gas emissions. Some methanotrophs are also metabolically flexible, growing at atmospheric methane concentrations by co-oxidizing hydrogen and carbon monoxide, which could allow them to persist in air independently of local methane supply ^25^. Methane concentrations vary predictably across ecosystem types, reflecting the uneven distribution of methane-emitting environments, both naturally occurring and anthropogenic, such as wetlands, rice fields, ruminant digestive systems, and landfills ^26^. The first step of methane oxidation is catalyzed by methane monooxygenase (MMO), which exists in particulate (pMMO) and soluble (sMMO) forms. The pMMO, encoded by *pmoCAB*, is more broadly distributed, and *pmoA* is widely used as a phylogenetic and functional marker for aerobic methanotrophs ^27,28^. Because methanotrophy is restricted to defined bacterial lineages and associated with characteristic MMO genes, detection of *pmoA* genes in aerosol metagenomes provides a specific indicator of methanotroph representation. Under a passive dispersal model, methane oxidation gene representation in aerosols should reflect source community composition proportionally and show no systematic variation across sites with different ambient methane concentrations. Under a context-dependent model, their representation in aerosols should instead vary with ambient methane availability, being more abundant where local methane is elevated.

Here, we test these alternative models across sites with contrasting ambient methane concentrations. We quantified airborne cell concentrations by flow cytometry and used shotgun metagenomics to compare taxonomic composition and functional gene repertoires between aerosols and paired source communities, focusing on methane monooxygenase genes (*pmoCAB* and the sMMO gene cluster) and extending the framework to broader C1 and air-relevant biogeochemical functions.

## Materials and Methods

### Study design and sample collection

Environmental samples were collected from seven sites across the lower mainland of British Columbia, Canada, to compare aerosol communities with paired underlying soil or water (Fig. 1, Table 1). Five focal sites represented ecosystem types expected to differ in ambient methane concentration (three wetlands and two anthropogenic sources). At two additional urban sites, aerosol DNA was insufficient for library construction, so these sites contributed flow cytometry data only. At each site, aerosols were collected as nine independent impinger collections of 4.5 m³ air each, and underlying environments were sampled as two soil replicates (or two water replicates at Burns Bog). Because DNA yields from individual aerosol collections were extremely low, aerosol collections from each site (eight or nine collections) were pooled prior to DNA extraction, yielding a single composite aerosol sample for shotgun metagenomic library preparation. Consequently, each site contributed one aerosol metagenome and two paired environmental metagenomes. As in many aerosol shotgun metagenomic studies, biological replication of aerosol samples was constrained by the low biomass typical of atmospheric collections, which often limits independent library preparation ^20^.

**Figure 1.**
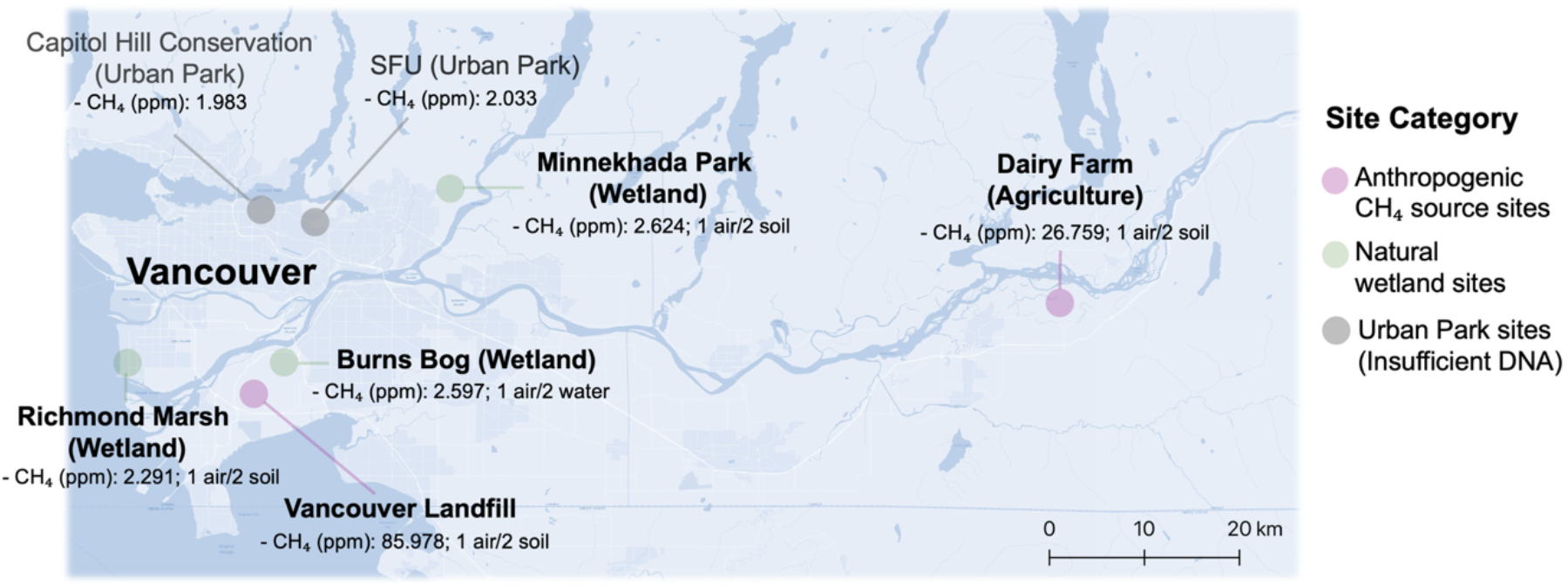
Sampling sites. Locations of the seven sampling sites, site categories, and ambient CH₄ concentrations measured during aerosol sampling. Site abbreviations are defined in Table 1.

**Table 1.** Sampling locations, environmental conditions, and samples collected and sequenced. Temperature, wind speed, and relative humidity (RH) are mean values during aerosol sampling, except at Minnekhada Park (*), where a single measurement was recorded. The VL methane concentration exceeded the instrument’s upper calibration limit (30 ppm) and is a lower bound. One composite aerosol sample (pooled from eight or nine collections) and two paired underlying environmental samples (soil or water) per site were processed for metagenomic sequencing. Aerosol libraries were constructed for five of seven sites.

| Site | Coordinates | CH<sub>4</sub> (ppm) | Temp. (°C) | Wind (m s<sup>-1</sup>) | RH (%) | Samples collected | Samples submitted for sequencing | Aerosol metagenome recovered |
| --- | --- | --- | --- | --- | --- | --- | --- | --- |
| Simon Fraser University (SFU) | 49°16'43"N<br>122°55'04"W | 2.03 | 8.6 | 1.01 | 75 | 9 Air/2 Soil | 1 Air/2 Soil | No (insufficient DNA) |
| Capitol Hill Conservation Area (CH) | 49°17'32"N<br>122°59'14"W | 1.98 | 9.2 | 0.59 | 76.5 | 9 Air/2 Soil | 1 Air/2 Soil | No (insufficient DNA) |
| Burns Bog (BB) | 49°07'08.9"N<br>122°59'42.3"W | 2.60 | 15.8 | 0.57 | 55.1 | 8 Air/2 Water | 1 Air/2 Water | Yes |
| Minnekhada Park (MK) | 49°18'07"N<br>122°42'01"W | 2.62 | 2.6* | 0* | 77.2* | 9 Air/2 Soil | 1 Air/2 Soil | Yes |
| Vancouver Landfill (VL) | 49°06'10"N<br>122°59'40"W | ≥ 86 | 3.4 | 0.8 | 87.8 | 8 Air/2 Soil | 1 Air/2 Soil | Yes |
| Richmond Marsh (RM) | 49°07'51"N<br>123°11'43"W | 2.29 | 12.8 | 0.42 | 70.2 | 9 Air/2 Soil | 1 Air/2 Soil | Yes |
| Dairy Farm (DF) | 49°11'30"N<br>121°49'18"W | 26.76 | 12.2 | 0.11 | 73.3 | 9 Air/2 Soil | 1 Air/2 Soil | Yes |

### Methane and meteorological measurement

Bulk air was collected into 10 L Tedlar® SCV gas sampling bags with Thermogreen® LB-2 septa (MilliporeSigma, Burlington, MA, USA) using a gas-tight syringe. Methane concentrations were measured within 24 h using cavity ring-down spectroscopy (Picarro G2210-i, Picarro Inc., Santa Clara, CA, USA) via a small sample introduction module (SSIM2). For each site, three replicate injections were performed from the same bag, and the mean value was used as the methane concentration at the time of aerosol sampling, as reported in Table 1. The VL sample exceeded the instrument’s upper calibration limit (30 ppm) and was not diluted before measurement. Because the instrument response is not linear above this limit, the true concentration may be higher, and the measured value is reported as a lower bound. Air temperature, relative humidity, and wind speed were measured at each site during aerosol sampling using a Kestrel 5000 weather meter (Nielsen-Kellerman, Boothwyn, PA, USA) and averaged over the sampling period (Table 1).

### Aerosol collection

Aerosols were collected using a liquid microimpinger (Coriolis μ, Bertin Technologies) ^29–31^ at 300 L min⁻¹ for 15 min (4.5 m³ air per collection), positioned 1.5 m above the ground to capture aerosols directly above each source environment. At this height, samples represent near-source air rather than well-mixed regional air ^32^. Sterile phosphate-buffered saline (PBS; pH 7.4, 15 mL) served as the collection medium. Liquid fractions were transported to the lab on ice and pooled across the eight or nine collections per site prior to filtration onto 25 mm, 0.22 μm polyethersulfone (PES) membrane filters (MilliporeSigma) for DNA extraction (N.M., J.P., M. Cleveland-Iliffe, N.B.D. and S.J.F., manuscript submitted). Field negative controls (PBS-filled cones mounted on the sampler for 15 min without airflow) were processed alongside samples at each site.

### Flow cytometry

Total and intact bacterial cell concentrations were quantified from four of the nine collections per site using a BD Accuri C6 Plus flow cytometer (BD Biosciences, San Jose, CA, USA), following previously established flow cytometry protocols for bacterial cell enumeration and membrane integrity assessment ^33,34^. Samples were stained with SYBR Green I to enumerate total cells and with SYBR Green I in combination with propidium iodide to differentiate intact cells from membrane-compromised cells ^35^. Each collection was measured in technical triplicate by volume-based acquisition of 150 µL. Field negative controls (two per site) were analyzed identically. Cell concentrations in air were calculated as C_air = (N / V_a) × V_L / V_air, where C_air is the cell concentration (cells m⁻³), N is the number of events counted, V_a is the analyzed volume (150 µL), V_L is the liquid volume recovered from the collection cone after sampling (12 to 13 mL, reduced from 15 mL by evaporation) and V_air is the air volume sampled (4.5 m³). This calculation assumes a collection efficiency of 1 and uniform aerosol concentration during each 15 min collection. Total cells in each pooled sample were estimated as site mean TCC multiplied by the total air volume pooled.

### DNA extraction and quantification

DNA was extracted from soil samples using the E.Z.N.A. Soil DNA Kit (Omega Bio-tek, Norcross, GA, USA) and from aerosol and water samples using the DNeasy PowerWater Kit (Qiagen, Hilden, Germany), processing entire filters. Field negative controls and extraction blanks were included to monitor contamination. DNA was quantified using the Qubit dsDNA High Sensitivity assay (Thermo Fisher Scientific, Waltham, MA, USA). DNA concentrations in all field negative controls and extraction blanks were below the detection limit of the Qubit assay, and library construction from field negative controls was unsuccessful.

### Metagenomic sequencing

Shotgun metagenomic libraries were prepared at the BC Cancer Genome Sciences Centre (Vancouver, BC, Canada) using a low-input, PCR-amplified library preparation protocol and sequenced on an Illumina NovaSeq 6000 platform (paired-end 150 bp), targeting ∼100 million read pairs per sample. Field negative controls did not yield usable libraries. In total, 15 libraries were retained for downstream analyses: five composite aerosol metagenomes and ten paired environmental metagenomes (two per site). Soil metagenomes from SFU and CH were excluded from analysis because no paired aerosol metagenome was available. Associations between aerosol cell counts, DNA yield, and sequencing depth were assessed using Spearman rank correlation. SFU DNA yield, below detection, was assigned 0.1 ng, which does not affect rank-based correlation.

### Metagenomic read processing and quality control

All bioinformatic analyses were performed on the Digital Research Alliance of Canada high-performance computing cluster (Fir). Raw paired-end reads were quality-assessed using FastQC ^36^ and processed with KneadData (v0.12.0) ^37^ for adapter trimming and quality filtering (Trimmomatic-based; sliding window 4:20, minimum length 50 bp). Post-filtering quality metrics were evaluated with FastQC and summarized across samples using MultiQC ^38^. All downstream analyses and visualizations were performed in R (v4.5.0) ^39^.

### Taxonomic profiling and diversity analyses

Quality-controlled paired-end reads were taxonomically classified using Kraken2 ^40^ with the PlusPF database (standard database plus protozoa and fungi; downloaded August 2025), and genus-level abundance estimates were refined using Bracken ^41^ at a read length of 150 bp. Downstream analyses were conducted at the genus level. Bacteria, Archaea, Eukaryota, and Viruses were analyzed separately to account for domain-specific sequencing depth and compositional structure. Eukaryotic profiles were restricted to fungi and protozoa represented in the reference database. Minimum relative abundance thresholds were applied prior to diversity analyses to remove classification noise (≥0.01% for Bacteria and Archaea, ≥0.005% for Eukaryota, and ≥0.001% for Viruses in at least one sample).

For alpha- and beta-diversity analyses, genus-level count tables were rarefied to the minimum library size observed within each domain (Bacteria: 416,709 reads; Archaea: 965 reads; Eukaryota: 63,349 reads; Viruses: 511 reads). Shannon diversity was calculated using phyloseq ^42^ and compared between sample types using Welch’s t-test. Community dissimilarity was quantified using Bray–Curtis distances ^43^ and visualized by non-metric multidimensional scaling (NMDS) ^44^. Differences in community composition between aerosol and paired environmental samples were tested using PERMANOVA (adonis2, 9,999 permutations) ^45^, and homogeneity of multivariate dispersion was evaluated using betadisper ^46^. To assess concordance between bacterial and eukaryotic community structures across samples, Procrustes analysis was performed in vegan, with significance evaluated by PROTEST (999 permutations) ^47^; Procrustes residuals were compared between sample types using a Wilcoxon rank-sum test.

### Differential abundance analysis

To identify genera showing differences in relative abundance between aerosol and paired environmental samples, differential abundance analysis was performed separately for Bacteria and Eukaryota using DESeq2 (v1.50.2) ^48^ on non-rarefied genus-level count tables derived from Kraken2/Bracken classification. A site-blocked design (∼ Site + SampleType) was used to account for baseline community differences among sites while estimating the overall aerosol versus paired-environment effect across the dataset. Because aerosol samples were represented by a single composite library per site, differential abundance estimates reflect dataset-level contrasts conditioned on site rather than within-site replication. Genera with fewer than 10 total counts across all samples were excluded prior to analysis. Size factors were estimated using the positive counts method to accommodate sparse data. Log_2_ fold changes were shrunk using the apeglm method ^49^ to reduce instability associated with low-count genera. Statistical significance was defined as a Benjamini–Hochberg adjusted p-value < 0.05. Genera with adjusted *p* < 0.05 and |log₂ fold change| ≥ 2 were considered differentially abundant.

### Metagenome assembly and gene catalog construction

Quality-controlled reads were co-assembled separately for each site using MEGAHIT (v1.2.9) ^50^, with each site-level co-assembly comprising one composite aerosol sample and two paired environmental samples. Open reading frames (ORFs) were predicted from contigs ≥1,000 bp (4,621,648 contigs) using Prodigal (v2.6.3) ^51^ in metagenomic mode, yielding 16,088,825 ORFs. Predicted proteins from all five assemblies were clustered at 95% amino acid identity using CD-HIT (v4.8.1) ^52^, giving a non-redundant catalog of 15,177,915 genes.

### Gene quantification and functional annotation

Quality-controlled reads from each sample were mapped independently to the non-redundant gene catalog constructed from site-level co-assemblies using Salmon (v1.10.1) ^53^ in quasi-mapping mode, treating the gene catalog as a reference transcriptome to obtain gene-level abundance estimates, with sequence- and GC-bias correction enabled. Gene abundances were estimated as transcripts per million (TPM), which normalize for gene length and sequencing depth, and converted to proportions within each sample.

Representative protein sequences from the non-redundant gene catalog were functionally annotated using eggNOG-mapper (v2.1.12) ^54^, which searched the eggNOG 5.0 database with DIAMOND (v2.1.8) ^55^ as the alignment engine. KEGG Orthology (KO) identifiers were retained for annotations meeting eggnog-mapper default confidence thresholds. All predicted ORFs were annotated; annotations assigned to eukaryotic or viral lineages based on eggNOG taxonomic scope were excluded to restrict functional analyses to prokaryotic gene content.

### Gene-centric functional profiling using curated marker genes

Gene-level TPM values were summed by KEGG Orthology (KO) identifier within each sample. Marker genes for C1 and trace-gas metabolism, nitrogen, phosphorus and sulfur cycling, VOC degradation, and persistence-related traits were defined *a priori* from canonical pathways, independently of their detection in the dataset (Supplementary Table S1), and summed within categories where indicated. This approach infers community-level functional potential, not pathway completeness or activity. For visualization, abundances were standardized as within-site z-scores, and KOs not detected in any sample were omitted from heatmaps. For site-level comparisons, the two environmental replicates per site were averaged, and log₂(aerosol / paired environment) fold changes were calculated from proportional TPM values; no fold change was calculated where a gene was not detected in the aerosol sample. Because pmoCAB and amoCAB share KEGG Orthology identifiers (K10944–K10946), the taxonomic affiliation of genes assigned to these KOs was checked, and those detected in aerosol metagenomes were affiliated with methanotrophs.

## Results

### Aerosol microbial cell concentrations vary across sites

Microbial cell concentrations in aerosols were quantifiable across all seven sites but site means differed by nearly 900-fold, reflecting strong site-level differences in airborne microbial load (Fig. 2A). Ambient methane concentrations at the time of aerosol sampling ranged from near-background levels (∼2 ppm) at wetland and urban sites to substantially elevated concentrations at DF (26.76 ppm) and VL (≥86 ppm) (Table 1). Publicly available monitoring data for Burns Bog (BB) and Richmond Marsh (RM) indicate that near-surface methane mixing ratios at these sites typically remain between 2 and 4 ppm ^56^, consistent with the values we measured at each site (2.60 and 2.29 ppm, respectively). Site mean total cell counts (TCC) ranged from 1.7 × 10³ cells per m³ of air at SFU to 1.5 × 10⁶ cells m⁻³ at DF (Fig. 2A). TCC was also high at VL (2.5 × 10⁵ cells m⁻³), largely because one collection taken immediately after new waste was deposited reached 9.5 × 10⁵ cells m⁻³; the other three VL collections ranged from 1.1 to 1.7 × 10⁴ cells m⁻³. BB was intermediate (5.6 × 10⁴ cells m⁻³), whereas MK, CH, RM and SFU had the lowest concentrations (1.7 to 6.1 × 10³ cells m⁻³). The fraction of total cells that were measured to be intact (ICC/TCC) varied markedly among sites and was independent of absolute cell load; for example, DF had the highest TCC but one of the lowest intact fractions (Fig. 2B). The intact fraction was highest at MK (65%) and VL (42%), which also showed the greatest variability among collections (32–96% and 21–61%, respectively), intermediate at CH (21%) and SFU (17%), and lowest at DF, BB, and RM (5%, 5%, and 2%, respectively). Previously reported total bacterial concentrations in near-surface air over land typically exceed 10⁴ cells m⁻³, although no wetland measurements were available ^57^, and reached 8.4 × 10⁵ bacteria-like particles m⁻³ at an outdoor site ^58^. TCC at BB, VL and DF fell within or above this range, whereas TCC at SFU, CH, MK and RM fell below it.

**Figure 2.**
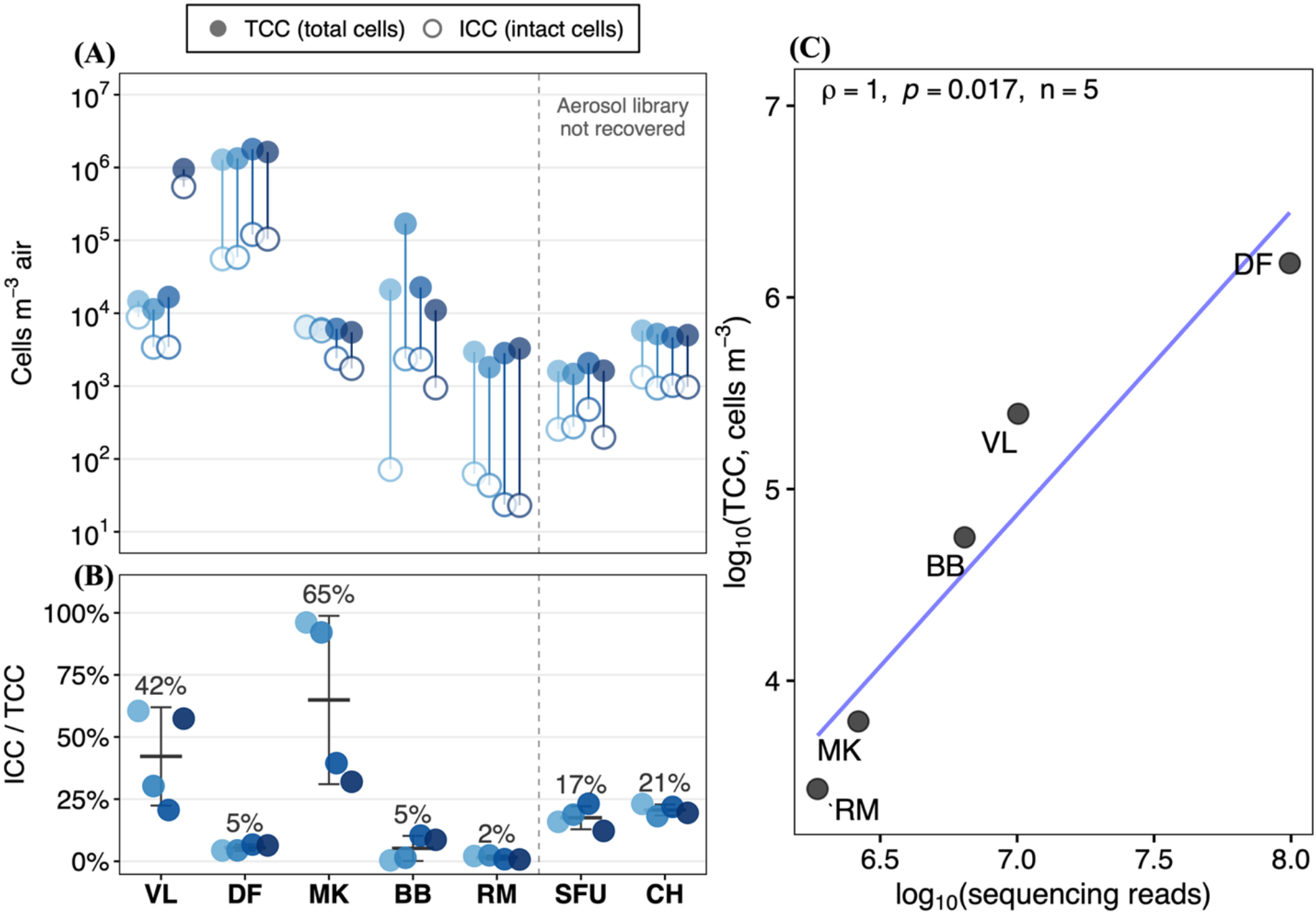
Aerosol cell concentrations quantified by flow cytometry. (A) Total (TCC; filled circles) and intact (ICC; open circles) cell count (cells m⁻³ air) for four of the nine impinger collections per site. Each point is the mean of technical triplicates; vertical segments link TCC and ICC from the same collection, and shades distinguish collections within each site. (B) Intact cell fraction (ICC/TCC) per collection, with site mean ± SD (n = 4); percentages indicate site means. Sites in A and B are ordered from high to low ambient CH₄ concentration. (C) Site mean TCC versus sequencing reads for the five aerosol metagenomes (log₁₀ scales), with Spearman ρ and *p* (n = 5); the line is for visualization only. Site abbreviations are defined in Table 1.

Field negative controls contained 0–359 cells m⁻³ (TCC) and 0–53 cells m⁻³ (ICC); ICC at RM (23–62 cells m⁻³) was close to its site controls, whereas all other values exceeded those of their site-specific controls. TCC was positively associated with sequencing depth across the five aerosol metagenomes (Spearman ρ = 1.00, *p* = 0.017, n = 5; Fig. 2C), although the small number of sites limits statistical power. Associations of ICC with sequencing depth and of TCC and ICC with DNA yield were positive but not significant (Supplementary Fig. 1). Pooled aerosol samples contained an estimated 6.9 × 10⁴ to 6.1 × 10⁷ cells (site mean TCC × air volume pooled). Libraries were obtained from all samples with ≥2.5 × 10⁵ cells, but outcomes overlapped below this level (RM succeeded with 1.1 × 10⁵ cells, whereas CH failed with 2.1 × 10⁵ cells). DNA yield separated outcomes more clearly: libraries were obtained from all samples with ≥6.75 ng DNA and from none with ≤4.83 ng (Supplementary Fig. 1).

### Domain composition of aerosol and paired environmental metagenomes

Bacteria accounted for most classified reads in all paired environmental samples, the underlying soil or water collected at the same site as each aerosol sample (96.9–99.6%) and in four of five aerosol samples (69.4–98.3%), whereas eukaryotic reads dominated the MK aerosol (79.5%), which also had the lowest DNA yield among the five aerosol libraries (6.75 ng). Overall, eukaryotic reads made up a larger and more variable share in aerosols (mean 26.0%, range 1.0–79.5%) than in paired environmental samples (mean 0.6%, range 0.1–2.5%; Supplementary Fig. 2, Supplementary Table S2). Archaeal and viral reads each accounted for less than 1% of classified reads in all samples.

### Bacterial and eukaryotic communities differ in aerosol-environment divergence

Bacterial and eukaryotic communities differed in the extent to which aerosol and corresponding environmental samples diverged compositionally (Fig. 3). Shannon diversity, which reflects both the number of taxa and how evenly reads are distributed among them, did not differ between aerosol and paired environmental samples for bacteria (Welch’s t-test, *p* = 0.156; Fig. 3A) but was higher in environmental samples than in aerosols for eukaryotes (*p* = 0.020; Fig. 3C). Aerosol eukaryotic communities were therefore less diverse than their source communities, whereas aerosol bacterial communities were not.

**Figure 3.**
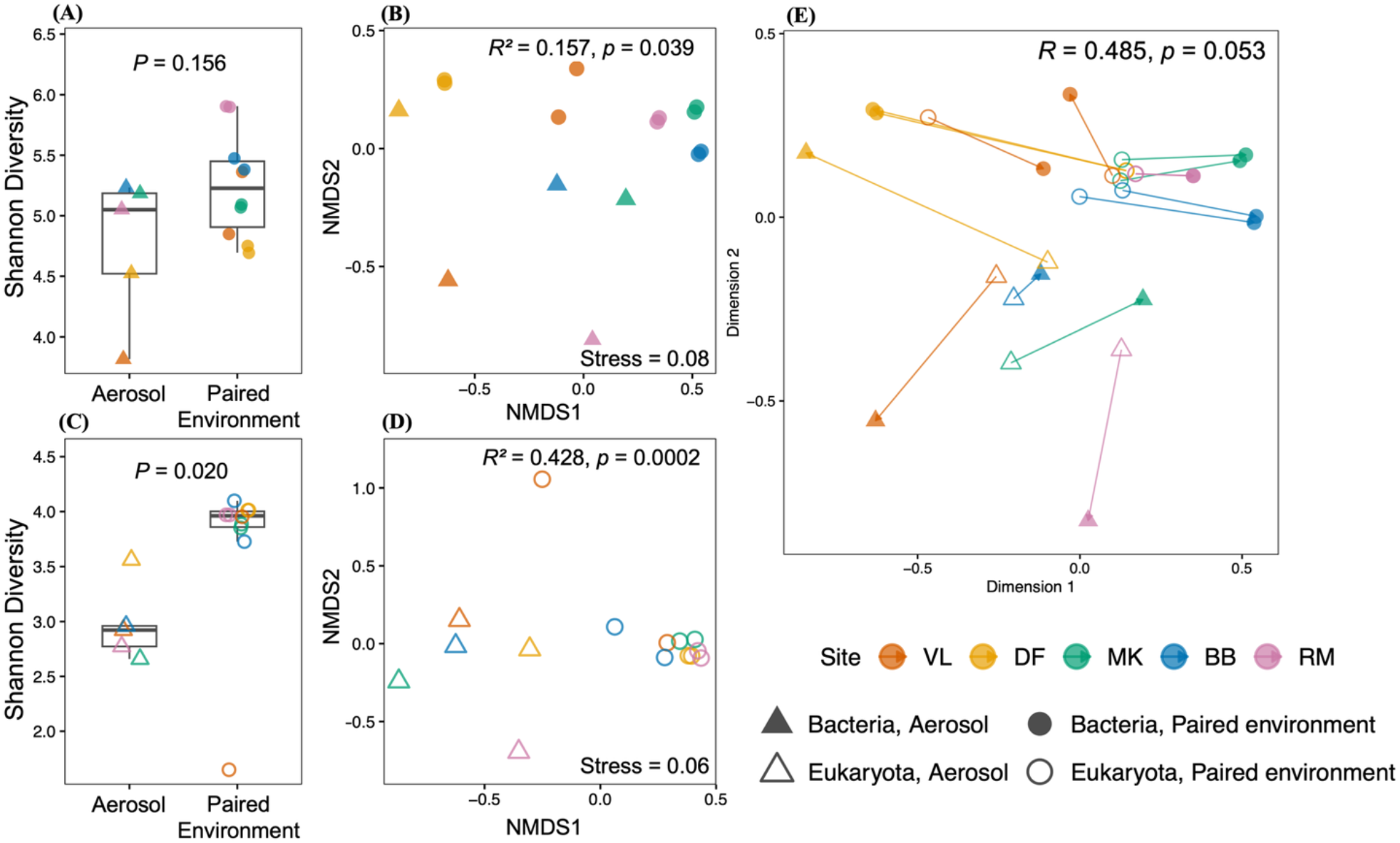
Bacterial and eukaryotic community diversity and composition in aerosol and paired environmental samples. Shannon diversity index (A, C; higher values indicate more taxa and more even abundances) and non-metric multidimensional scaling (NMDS) ordination based on Bray–Curtis dissimilarity (B, D; axes are unitless, and closer points indicate more similar community composition) of bacterial (A, B) and eukaryotic (C, D) communities. (E) Procrustes analysis of bacterial and eukaryotic ordinations across all samples; arrows connect the eukaryotic to the bacterial position of each sample, with longer arrows indicating a greater mismatch between bacterial and eukaryotic community patterns. Colours denote site; triangles denote aerosol and circles paired environmental (soil or water) samples; filled and open symbols denote bacteria and eukaryotes, respectively. Boxplots show medians and interquartile ranges. Welch’s *t*-test *p* (A, C), PERMANOVA *R*² and *p* (9,999 permutations) with NMDS stress (B, D; values below 0.2 indicate an adequate two-dimensional representation) , and PROTEST *r* and *p* (E) are shown in each panel. Analyses were performed at the genus level on rarefied counts (A–D) or relative abundances of non-rarefied counts (E). Aerosol, n = 5; paired environmental, n = 10.

NMDS ordination of bacterial communities based on Bray–Curtis dissimilarity showed partial overlap between aerosol and paired environmental samples (Fig. 3B), with sample type explaining 15.7% of compositional variance (PERMANOVA pseudo-F(1,13) = 2.42, R² = 0.157, *p* = 0.039). Within-group dispersion, which can inflate PERMANOVA results when it differs between groups, did not differ between sample types (betadisper F(1,13) = 0.34, *p* = 0.574), indicating that the PERMANOVA result reflects a difference in composition rather than in variability. Bacterial aerosol samples were broadly distributed across ordination space and clustered closer to their paired environmental samples than to aerosols from other sites. Eukaryotic communities, on the other hand, showed clear separation between aerosol and paired environmental samples, with sample type explaining 42.8% of compositional variance (PERMANOVA pseudo-F(1,13) = 9.74, R² = 0.428, *p* = 0.0002), with no evidence of differential dispersion (betadisper F(1,13) = 0.18, *p* = 0.741). Eukaryotic aerosol communities from different sites were more similar to one another than to their paired environmental samples (Fig. 3D). Thus, aerosol bacterial communities retained the composition of their local sources, whereas aerosol eukaryotic communities shared a similar composition across sites.

Procrustes analysis tested whether samples with similar bacterial communities also had similar eukaryotic communities. This correspondence was moderate and not significant (PROTEST r = 0.485, *p* = 0.053; Fig. 3E), and the mismatch between bacterial and eukaryotic patterns in individual samples (Procrustes residuals) did not differ between aerosol and paired environmental samples (Wilcoxon rank-sum test, *p* = 0.51). Archaeal and viral communities showed no significant differences between aerosol and paired environmental samples in either diversity or composition (PERMANOVA *p* > 0.05 for all comparisons; Supplementary Fig. 3) and, given their low read share, were not examined further.

### A subset of genera is consistently enriched in aerosols

A limited subset of genera showed consistent differences in representation between aerosol and paired environmental samples after accounting for site-level variation (Fig. 4). Most genera did not differ significantly after accounting for site effects (89.4% of bacterial and 92% of eukaryotic genera). Of those that did, 141 bacterial and 5 eukaryotic genera were enriched in aerosols, and 7 bacterial and 2 eukaryotic genera were depleted in aerosols relative to paired environmental samples (Supplementary Table S3). Bacterial genera with higher relative abundance in aerosols than in environmental samples included *Staphylococcus*, *Mammaliicoccus*, *Sellimonas*, *Jeotgalicoccus*, *Ligilactobacillus*, *Weissella*, *Aerococcus*, *Nosocomiicoccus*, and *Ruoffia* (Fig. 4A). Among eukaryotes, aerosol-enriched genera were exclusively fungal, including *Marasmius*, *Psilocybe*, *Rhizoctonia*, *Botrytis*, and *Debaryomyces* (Fig. 4B). Genus-level composition across all four microbial domains (Supplementary Fig. 4) further contextualized these patterns: bacterial aerosol communities were broadly dispersed across many low-abundance genera, while eukaryotic aerosols were comparatively enriched in spore-forming and filamentous fungal taxa.

**Figure 4.**
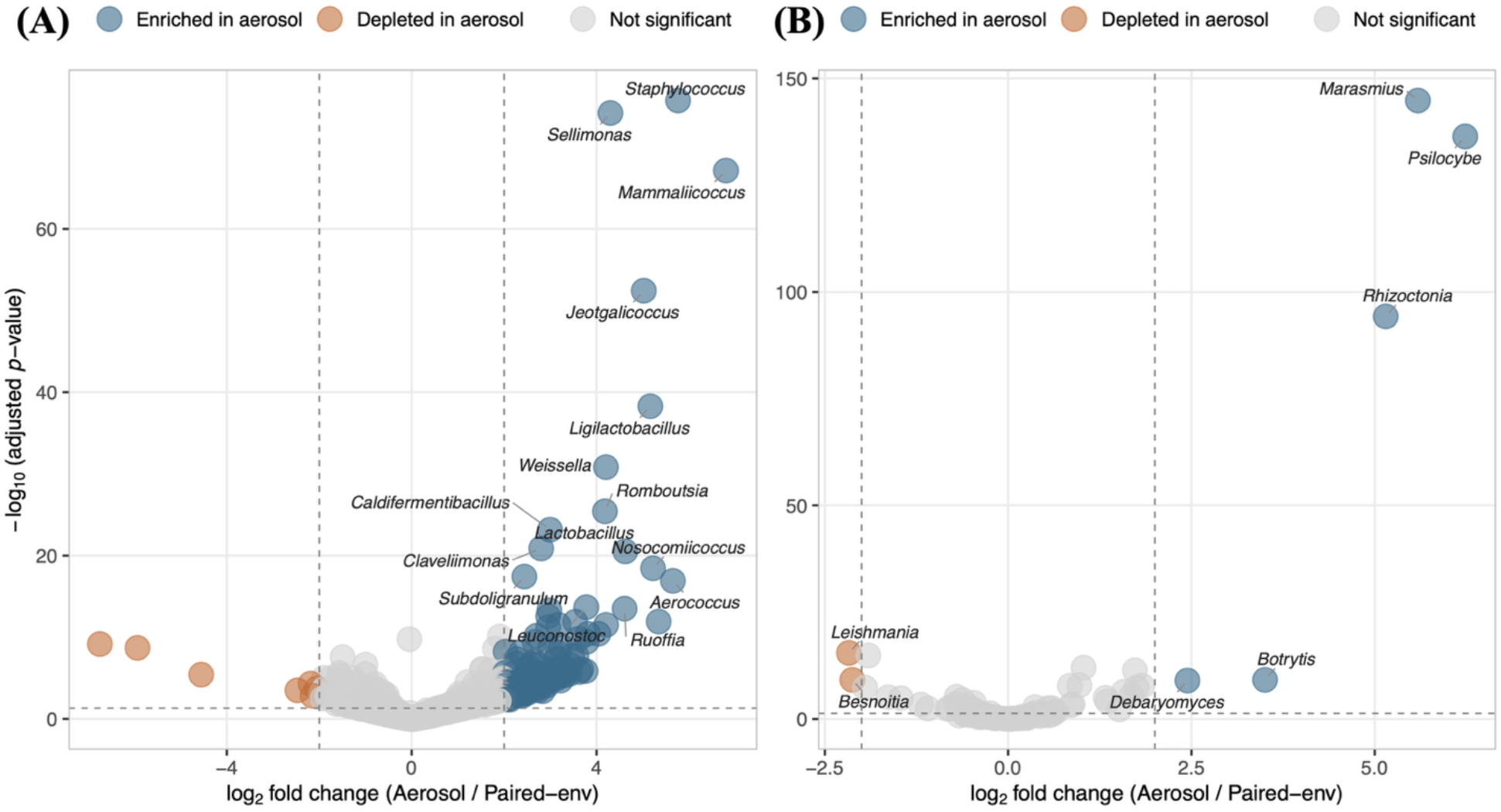
Differentially abundant bacterial and eukaryotic genera between aerosol and paired environmental samples. Differential abundance of (A) bacterial and (B) eukaryotic genera between aerosols (n = 5) and paired environmental samples (n = 10) (Kraken2/Bracken). Points represent individual genera plotted by log_2_ fold change (aerosol / paired-environment) on the x-axis and −log_10_ adjusted p-value on the y-axis (higher values indicate stronger statistical support). Differential abundance was assessed using DESeq2 with site-blocked design to account for site-level community variation, with Benjamini–Hochberg correction for multiple testing. Genera with |log_2_ fold change| ≥2 and adjusted *p* < 0.05 were considered significant. Genera enriched in aerosols are shown in blue, those depleted in aerosols in orange, and non-significant genera in grey. The top 15 differentially abundant taxa ranked by adjusted p-value were labelled.

### Methane oxidation gene representation in aerosols is consistent with local methane availability

Methane oxidation gene representation in aerosols differed among sites in line with ambient methane concentration (Fig. 5). Particulate methane monooxygenase genes (*pmoCAB*) were detected in aerosol metagenomes from VL (≥86 ppm CH₄) and DF (26.76 ppm CH₄), but were not detected in aerosol metagenomes at the three wetland sites (BB, MK, RM; ∼2 ppm CH₄). At VL and DF, the soluble methane monooxygenase subunit *mmoC* and methanol dehydrogenase markers (*xoxF*, *mxaI*) had higher relative abundance in aerosols than in paired environmental samples. Detection of *pmoCAB* genes in aerosols at VL and DF was consistent with the presence of canonical methanotrophic genera in read-based taxonomic profiles at these sites (Supplementary Fig. 5). Across sites, summed *pmoCAB* abundance was lower in aerosols than in paired environmental samples. However, the magnitude of depletion differed among sites: it was smallest at VL and DF, where ambient CH₄ concentrations were highest, whereas *<u>pmoCAB</u>* was not detected in aerosols at the wetland sites (Fig. 5B).

**Figure 5.**
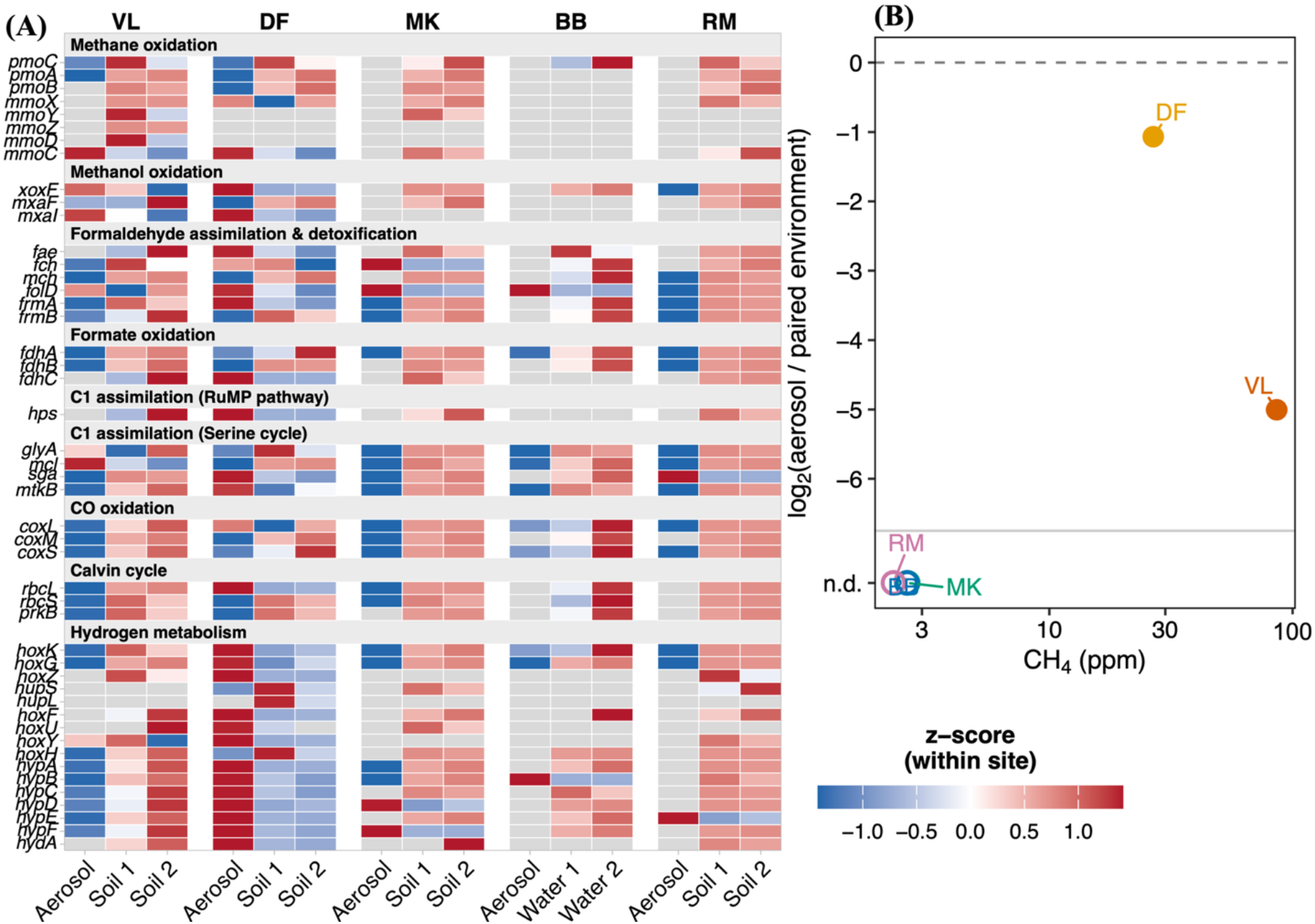
C1 metabolism and methane oxidation gene potential across aerosol and paired environmental samples. (A) Relative abundance of marker genes in nine functional categories of C1 and trace-gas metabolism, grouped under category headers. Gene abundances were derived from read recruitment to a non-redundant gene catalog, expressed as proportional TPM, and standardized as within-site z-scores for visualization. Grey tiles indicate no detection. Within each site, samples are ordered with aerosol first, followed by paired environmental samples (soil, or water at BB). (B, inset) log₂(aerosol / paired environment) of summed *pmoCAB* proportional TPM per site versus ambient CH₄ concentration (log₁₀ scale). Filled circles show sites where *pmoCAB* was detected in the aerosol metagenome; open circles (n.d.) show sites where it was not detected, for which no ratio could be calculated. Colours denote site.

Beyond methane oxidation, carbon monoxide oxidation genes (*coxLMS*) were detectable in aerosols at multiple sites, including wetland sites BB and RM. In contrast, most other marker genes associated with C1 processes (metabolism of one-carbon compounds such as methane, methanol and carbon monoxide), including those for formate oxidation (*fdhABC*), C1 assimilation via the RuMP pathway (*hps*), serine cycle (*glyA*, *mtkB*), and Calvin cycle (*rbcLS*, *prkB*), generally had higher relative abundance in paired environmental samples than in aerosols across sites. At wetland sites, these marker genes were frequently detected in paired environmental samples but were not detected or present at low levels in aerosols. Hydrogen metabolism genes exhibited site-specific patterns. At DF, membrane-bound hydrogenase subunits (*hoxK*, *hoxG*, *hoxZ*), soluble uptake hydrogenase components (*hupS*, *hupL*, *hupC*), and maturation factors (*hypA–F*) had higher relative abundance in aerosols than in paired environmental samples. *hoxG* and *hoxK* (large and small subunits of the membrane-bound [NiFe] uptake hydrogenases, respectively) were detectable in aerosol metagenomes across all five sites.

Assembly-based gene-centric profiling was extended to markers associated with volatile organic compound (VOC) and complex carbon degradation, nitrogen, phosphorus, and sulfur cycling, and survival-related traits across all five sites (Supplementary Figs. 6–10). Marker genes from each functional category were detected in aerosol metagenomes at one or more sites, confirming that the gene catalog captured functional signal across all categories. Relative abundance of these markers varied among sites, and no functional category showed a consistent directional shift between aerosol and paired environmental samples across all sites.

## Discussion

Methane oxidation gene representation in near-source aerosols was more consistent with local ambient methane concentration than with passive dispersal of source community functional potential. Methane oxidation genes had consistently lower relative abundance in aerosols than in paired source communities. Under passive dispersal, however, this environment-to-aerosol ratio would be similar across sites. Instead, *pmoCAB* genes were detectable in aerosol metagenomes only at VL and DF, where ambient methane concentrations exceeded 25 ppm, and were not detected at wetland sites where methane was near atmospheric background. This pattern suggests that the aerosol representation of specialist functional genes differs among sites in line with local substrate availability, rather than following a fixed environment-to-aerosol ratio. By contrast, nitrogen, phosphorus, sulfur, and VOC degradation markers showed no consistent directional shifts between aerosol and paired environmental samples across sites, providing no evidence that these functions are selectively enriched during aerosolization or atmospheric residence. More broadly, these findings are consistent with a context-dependent model of aerosol community assembly in which the representation of specialized metabolic genes reflects not only dispersal from a source environment, but also localized environmental conditions.

At the community level, the broad correspondence between aerosol and source community composition observed here is consistent with dispersal-based models in which airborne microbiomes largely reflect microbial inputs from local sources. For example, analyses of atmospheric aerosol samples show that many airborne bacteria can be traced to surrounding soils ^59^. However, the consistent enrichment of specific bacterial genera in aerosols indicates that aerosol composition is not a proportional sample of the underlying source, consistent with structured recurrence and diel patterns reported in long-term aeromicrobiome surveys ^9,60^. Functional metagenomic analyses of atmospheric aerosols and dust show that airborne gene repertoires can reflect regional source inputs ^1,61^. Our findings extend this by suggesting that the representation of specific functional genes, i.e., methane oxidation genes, varies with the local availability of methane, the growth substrate of methanotrophs, rather than simply tracking overall source community composition. In addition to the detection of *pmoCAB*, *mmoC*, which encodes the soluble methane monooxygenase reductase, had higher relative abundance in aerosols than in paired environmental samples at VL and DF, the sites with the highest ambient methane concentrations. Because *mmoC* homologues occur in multiple monooxygenase systems, its detection alone does not confirm the presence of soluble methane monooxygenase. Nevertheless, its co-occurrence with *pmoCAB* at methane-rich sites, and lack of detection at near-background sites is consistent with a methanotrophic source population whose aerosol representation varies with ambient methane availability.

The detection of methane oxidation genes at methane-rich sites further raises the question of whether trace-gas metabolism could support microbial persistence during atmospheric transport under organic carbon limitation. Laboratory studies show that some methanotrophs can oxidize trace concentrations of CH_4_, and in some cases can sustain growth on air by co-oxidizing atmospheric CH_4_ together with CO and H_2_ ^62^. Air-associated methanotrophs may also remain metabolically responsive when supplied with methane under aerosolized conditions in controlled lab experiments; for example, rotating gas-phase bioreactor experiments demonstrated ¹³C-methane incorporation into DNA in airborne methanotrophic enrichments ^63^, though these experiments used methane concentrations far exceeding ambient atmospheric levels, limiting direct comparison to natural aerosol conditions. These observations are consistent with growing evidence that atmospheric trace-gas oxidation can support microbial persistence in energy-limited environments, including oligotrophic cave ecosystems where trace-gas oxidation supports community-level primary production ^64^, and with the broader framework of atmospheric trace-gas metabolism ^15,65^. In this dataset, methane monooxygenase genes were restricted to the two methane-rich sites, whereas other trace-gas markers such as CO oxidation (*coxLMS*) and hydrogenase genes were detectable across more sites. This contrast suggests that methane oxidation potential in aerosols depends on local methane availability. The broader detection of CO and H_2_ oxidation genes may reflect their wide distribution across source communities ^66^. Alternatively, because CO and H_2_ were not measured, the representation of these genes may similarly track local concentrations of these gases. These possibilities remain open because shotgun metagenomes resolve genetic potential rather than activity. Establishing whether airborne methanotrophs are metabolically active, and by which mechanism, requires activity-resolved approaches such as metatranscriptomics, stable isotope labelling or controlled aerosolization experiments. What the present data establish is the prior step: local environmental conditions shape which functional genes are present in near-source aerosols.

Eukaryotic communities diverged more strongly between aerosols and paired environments than bacterial communities, with fungal genera particularly enriched in aerosols. This divergence need not reflect stronger selection during aerosolization. Several enriched genera, including the mushroom-forming *Marasmius* and *Psilocybe* and the plant pathogen *Botrytis*, release spores from aboveground fruiting bodies and plant tissue that our paired soil and water samples did not capture, and active spore release can decouple airborne fungal composition from fungal abundance in source habitats ^67,68^. Bacteria and fungi may also differ in the spatial and temporal dynamics of their dispersal ^69^. In addition, because eukaryotic genomes are typically much larger than bacterial genomes, eukaryotic read shares likely overstate their cellular abundance, particularly in low-biomass samples. This may contribute to the eukaryote dominance in the MK aerosol, which had the lowest DNA yield among sequenced aerosols, although a single composite sample per site cannot resolve the cause. Among bacteria, several bacterial genera enriched in aerosols in this study, including *Staphylococcus*, *Mammaliicoccus*, and *Ligilactobacillus*, are characteristic of animal-associated microbiomes and have been detected in airborne emissions from livestock facilities ^70,71^, suggesting that nearby animal activity contributed to local aerosol bacterial composition at some sites. More broadly, bacterial enrichment in aerosols may reflect contributions from local sources not captured by the paired samples, such as livestock at DF, preferential aerosolization of some taxa, or both.

Together, these results show that aerosol microbiomes are not proportional subsets of underlying source communities: bacterial aerosol communities largely mirrored local sources, eukaryotic communities diverged in ways consistent with dispersal traits and unsampled aboveground sources, and methane oxidation gene representation was consistent with local methane availability. These conclusions are constrained by low aerosol biomass, which required pooling of collections and precluded within-site replication, and by metagenomes capturing functional potential rather than activity. Replicated aerosol sampling across broader methane gradients, combined with activity-resolved approaches noted above, will be needed to test whether these signatures reflect metabolically active populations; the cell and DNA yields that supported library construction here (∼2.5 × 10⁵ cells or ∼7 ng DNA; Fig. 2C, Supplementary Fig. 1) provide a practical benchmark for planning such sampling. More broadly, predicting the composition and functional potential of airborne microbiomes will require accounting for local environmental context and source-specific dispersal, not passive dispersal alone.

## Acknowledgements

This work was supported by the Government of Canada’s New Frontiers in Research Fund (NFRF), grant NFRFE-2022-00163. The authors wish to acknowledge Canada’s Michael Smith Genome Sciences Centre for metagenomic library preparation and sequencing. This research was enabled in part by support provided by the BC DRI Group and the Digital Research Alliance of Canada (alliancecan.ca). Claude AI (Anthropic) was used for language improvement of the manuscript.

## Author contribution

S.J.F. conceived and supervised the study and secured funding. N.B.D. and N.Z. contributed to sampling design and funding acquisition; N.B.D. also contributed to conceptualization and methodology. J.P. and N.M. conducted aerosol field sampling, with N.M. developing and optimizing the aerosol sampling methodology. N.M. performed flow cytometric enumeration and aerosol DNA extraction. N.M. and A.K. performed the CRDS methane measurements in N.Z.’s laboratory, with A.K. providing training. J.P. extracted DNA from paired environmental samples, performed the metagenomic and statistical analyses, prepared the figures, interpreted the data, and wrote the original manuscript. S.M.B.K. contributed to discussions on the analyses and interpretation of the results. All authors reviewed and edited the manuscript and approved the final version.

## Competing interest

The authors declare that they have no competing interests

## Funding

This work was supported by funding from the Government of Canada’s New Frontiers in Research Fund (NFRFE-2022-00163).

## Data availability

Raw metagenomic sequencing reads generated in this study have been deposited in the European Nucleotide Archive under project accession PRJEB109287 and will be released upon publication. Associated sample metadata are available within the same project record.

## Notes

### Competing Interest Statement

The authors have declared no competing interest.

