## Supplementary_Figures_Tables for "Methane oxidation genes are only present in bioaerosols at sites with high methane concentrations"

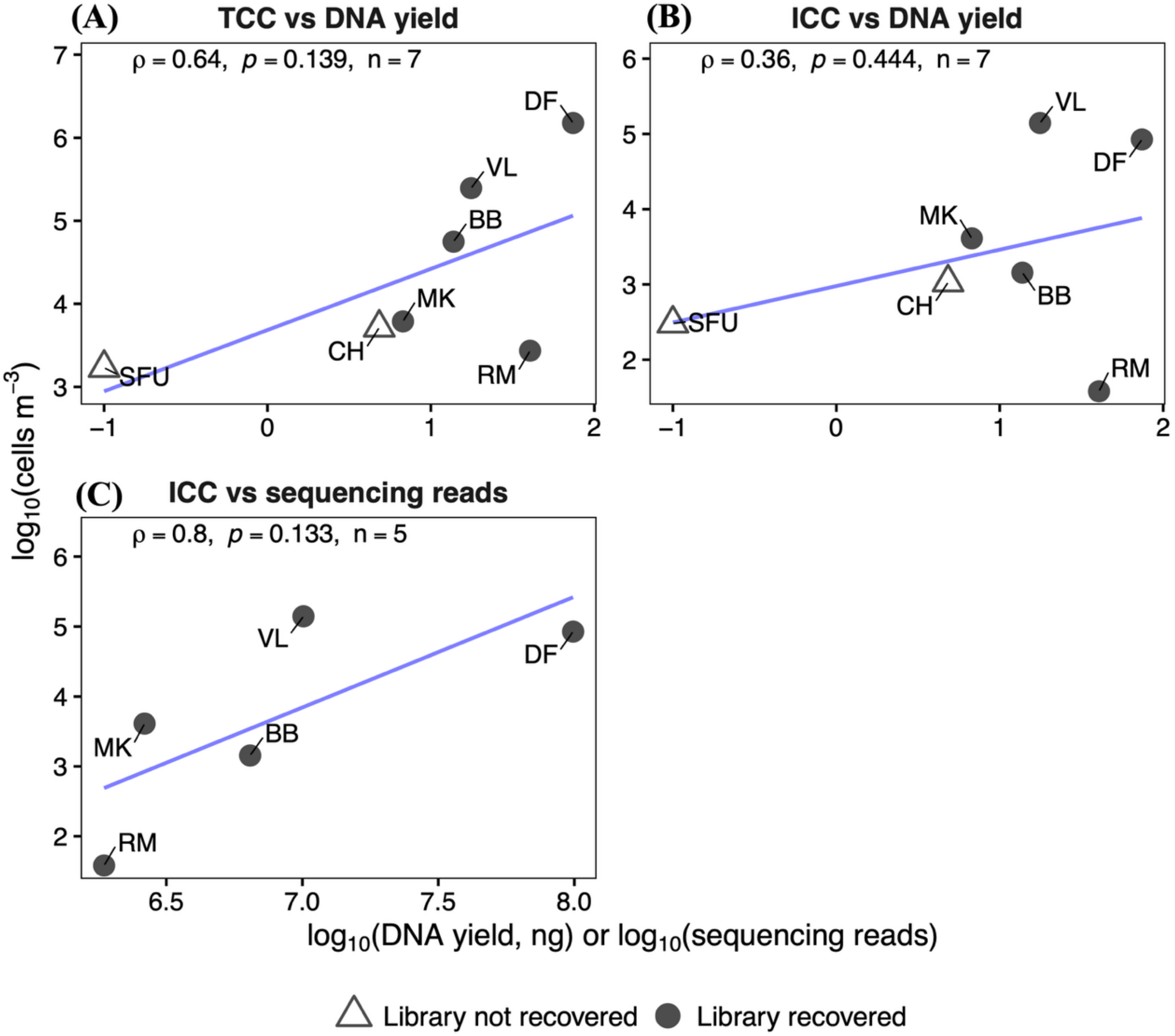


**Supplementary Figure 1 | Relationships between aerosol cell counts, DNA yield and sequencing depth.** Site mean TCC (A) and ICC (B, C) plotted against extracted aerosol DNA (A, B) and sequencing reads (C), on $\log_{10}$ scales. Open triangles indicate sites where aerosol libraries were not recovered (SFU, CH), shown in DNA panels only; SFU DNA, below detection, is plotted at 0.1 ng. Spearman ρ, *p*, and n are shown per panel; lines are for visualization only.

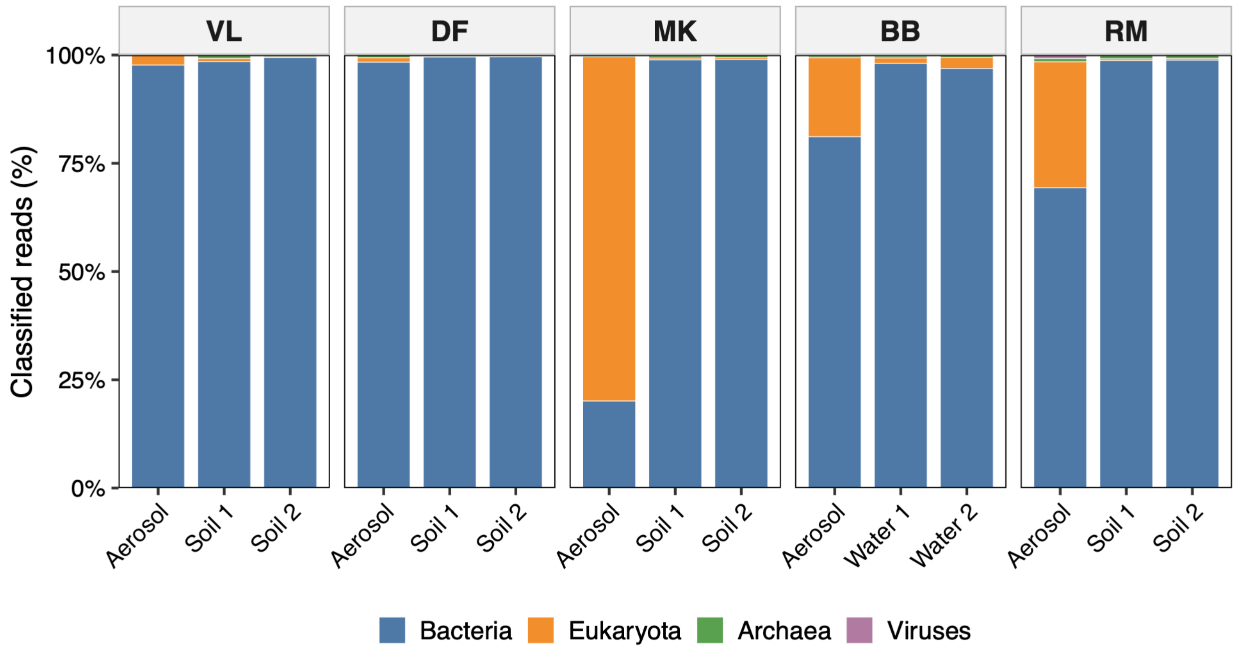


**Supplementary Figure 2 | Domain composition of aerosol and paired environmental metagenomes.** Percentage of classified reads assigned to Bacteria, Eukaryota, Archaea and Viruses in each aerosol and paired environmental (soil, or water at BB) sample (Kraken2/Bracken; human reads removed). Archaea and Viruses each accounted for less than 1% of reads in all samples and are barely visible; exact values are given in Supplementary Table S2. Sites are ordered from high to low ambient $CH_4$ concentration.

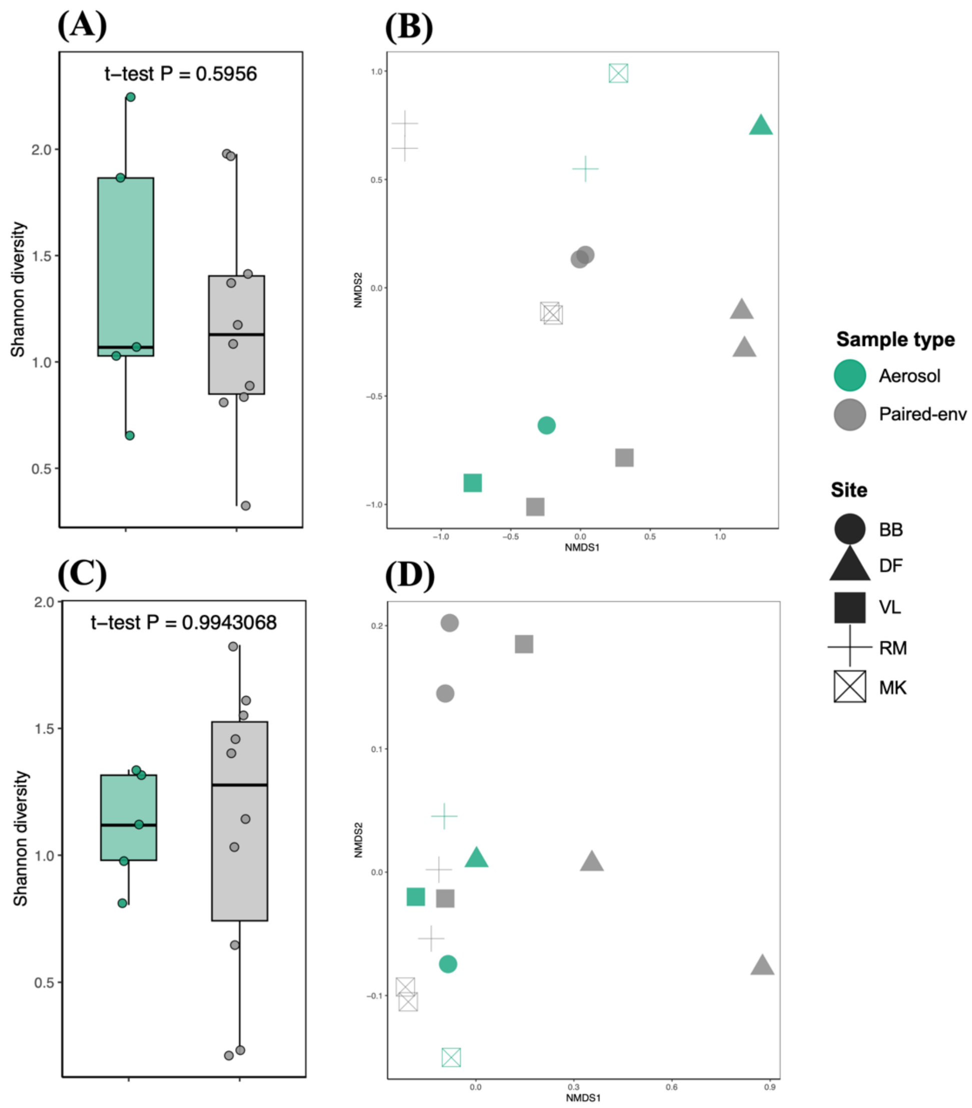


**Supplementary Figure 3 | Archaeal and viral community diversity in aerosol and paired environmental samples.** (A, C) Shannon diversity of archaeal (A) and viral (C) communities at the genus level (Kraken2/Bracken). Boxplots show medians and interquartile ranges; two-sample t-tests were used for group comparisons. Archaeal ($P = 0.596$) and viral ($P = 0.994$) Shannon diversity did not differ between aerosol ($n = 5$) and paired environmental samples ($n = 10$). (B, D) NMDS ordinations of archaeal (B) and viral (D) community composition based on Bray–Curtis dissimilarity. Colors denote sample type (green, aerosol; grey, paired environmental) and shapes denote site. Group differences were tested using PERMANOVA (adonis2, 9,999 permutations), with dispersion assessed by betadisper. Archaeal community composition showed no significant compartmental differentiation (PERMANOVA pseudo-$F(1,12) = 1.85$, $R^2 = 0.133$, $P = 0.074$; betadisper $F(1,12) = 0.019$, $P = 0.901$), and viral community composition did not differ between compartments (PERMANOVA pseudo-$F(1,12) = 0.99$, $R^2 = 0.076$, $P = 0.385$; betadisper $F(1,12) = 1.09$, $P = 0.363$). All analyses were performed at the genus level on rarefied count tables.

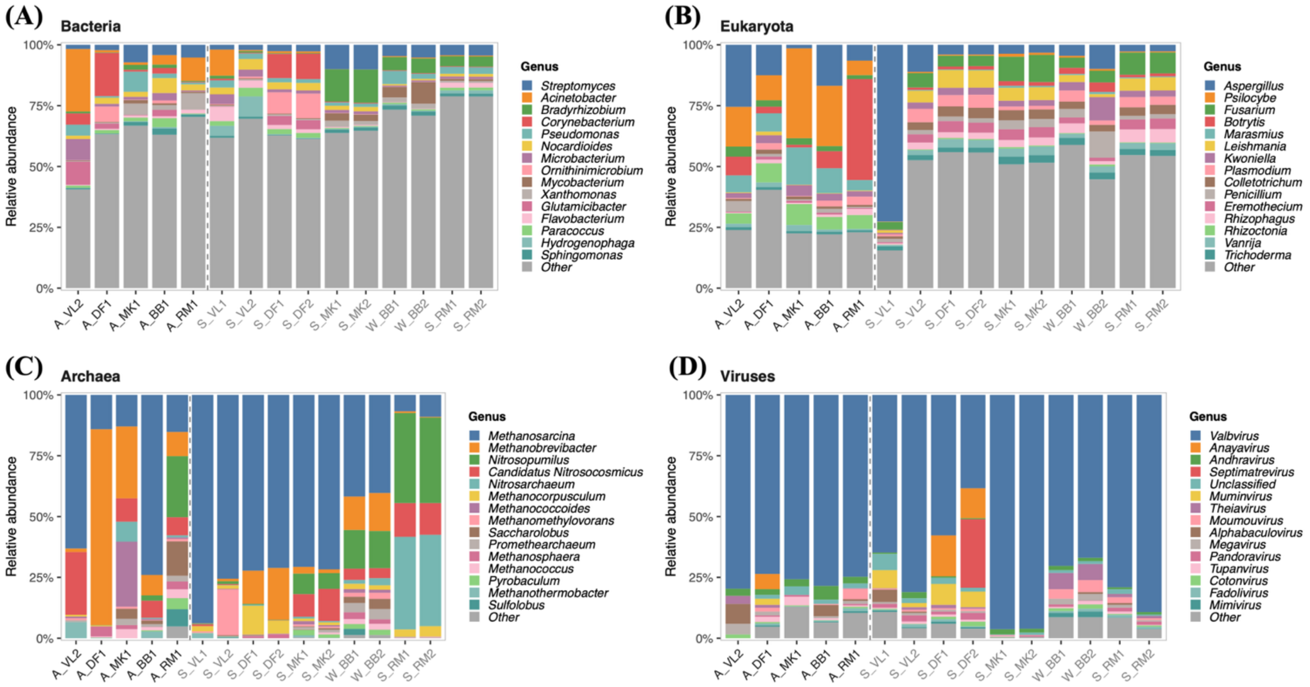


**Supplementary Figure 4 | Genus-level community composition across aerosol and paired environmental samples for all four microbial domains.** Stacked bar plots show the relative abundance of the top 15 genera per domain across all 15 samples. (A) Bacteria, (B) Eukaryota, (C) Archaea, and (D) Viruses. Samples are ordered with aerosol samples first (prefixed A_), followed by paired soil (prefixed S_) and water (prefixed W_) environmental samples with site names. Relative abundance was calculated from rarefied genus-level count tables (Kraken2/Bracken) and expressed as a proportion of total classified reads per sample within each domain. Genera not within the top 15 are collapsed into "Other." Grey dashed line separates aerosol from paired environmental samples.

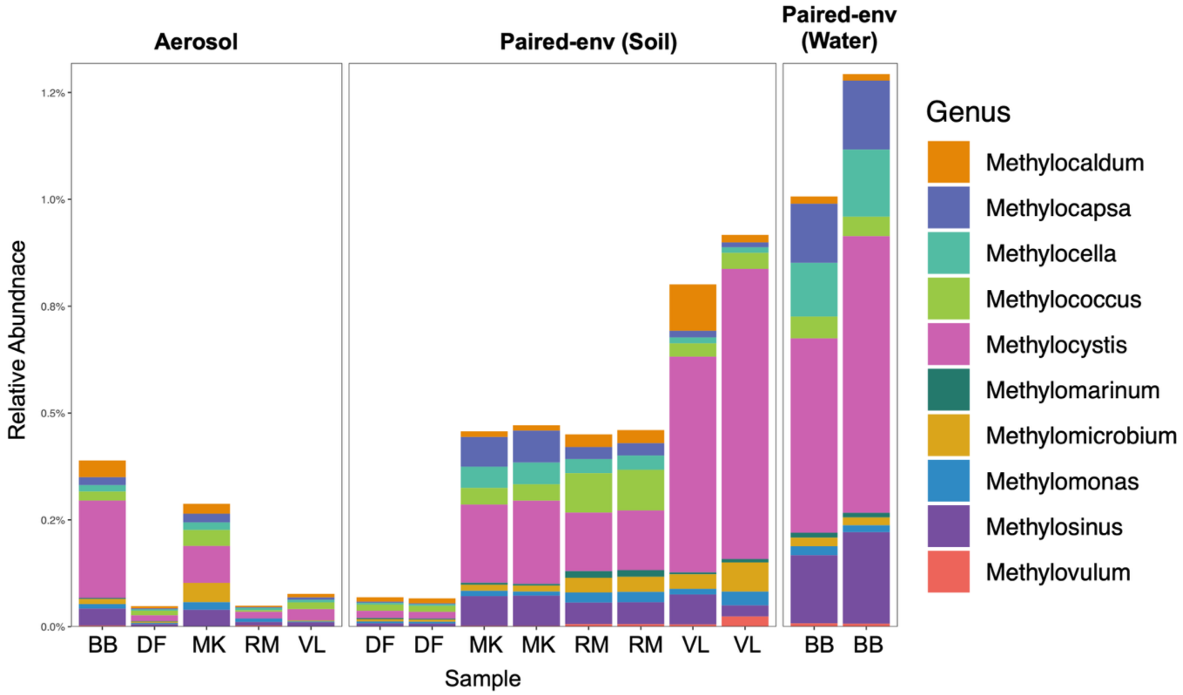


**Supplementary Figure 5 | Relative abundance of methanotrophic bacterial genera across aerosol and paired environmental samples.** Stacked bar plots show the relative abundance of recognized methanotrophic bacterial genera at the genus level (Kraken2/Bracken classification) across all samples. Bars represent individual samples grouped by site, including aerosols (n = 5) and paired soil or water samples (n = 10). Relative abundance is expressed as the percentage of total bacterial reads per sample. Methanotrophic taxa were detected across multiple sites but comprised a small fraction of total bacterial community composition.

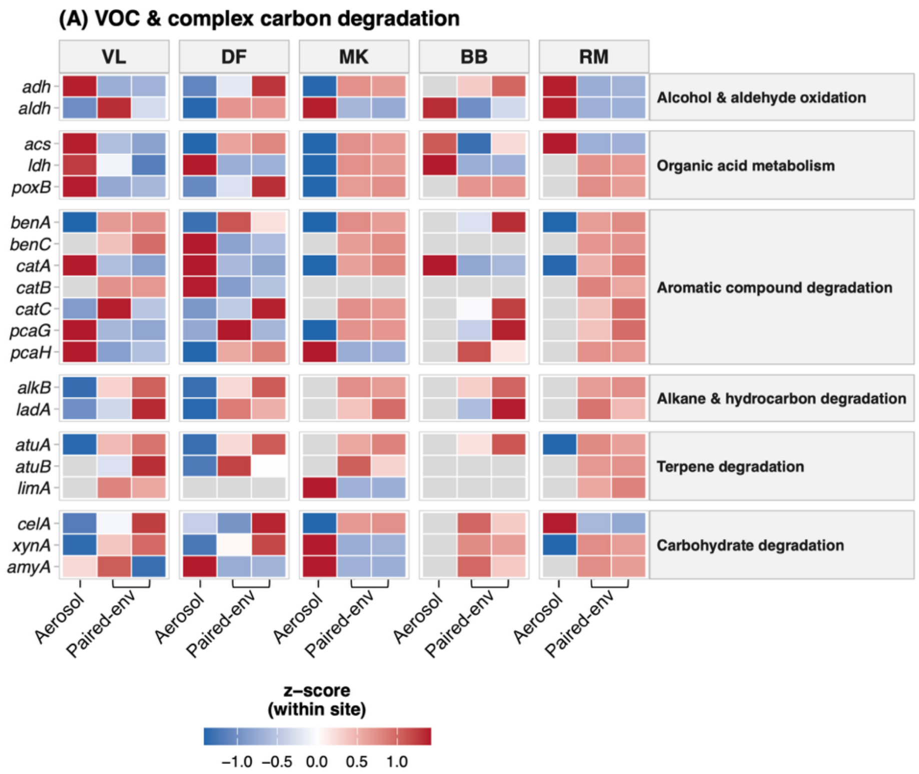


**Supplementary Figure 6 | VOC and complex carbon degradation gene potential across aerosol and paired environmental samples.** Gene abundances are z-score normalized within each site; grey tiles indicate absence of detection. See Fig. 5 for methodological details.

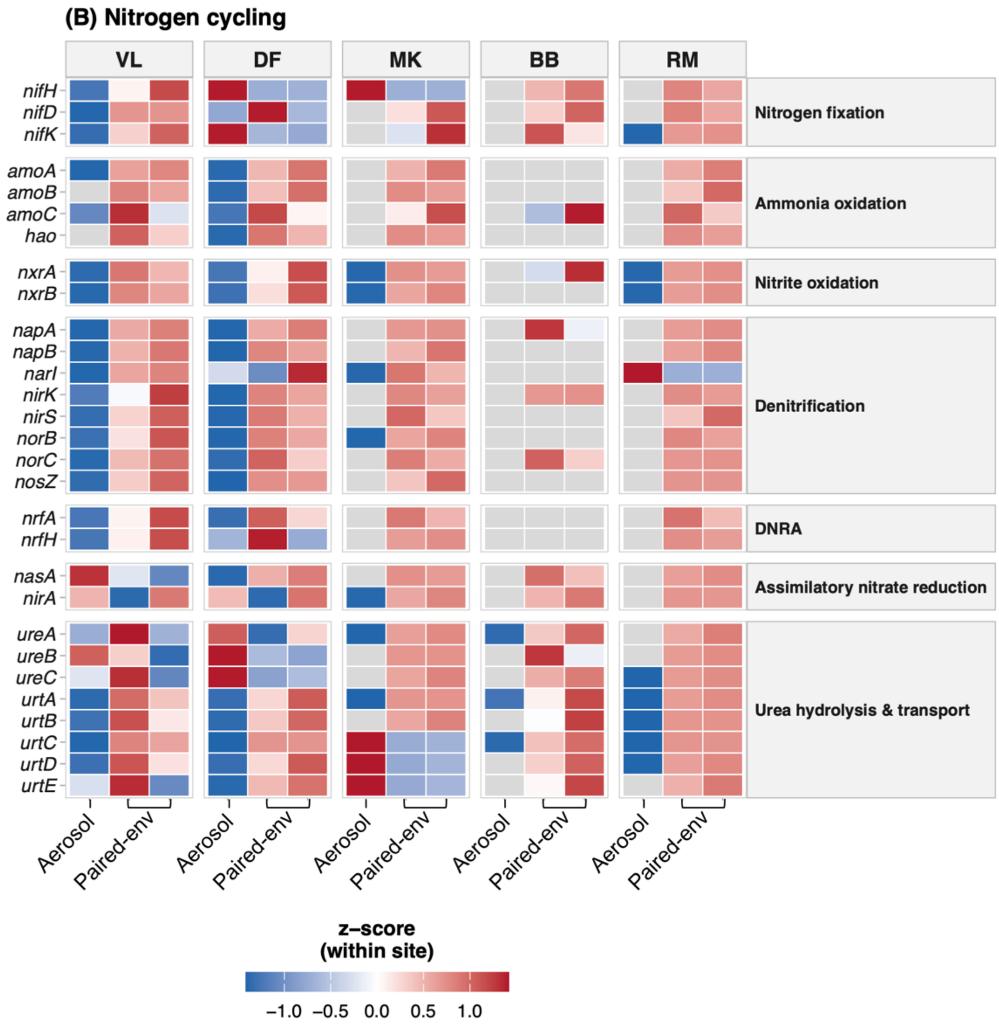


**Supplementary Figure 7 | Nitrogen cycling gene potential across aerosol and paired environmental samples.** As Supplementary Figure 5.

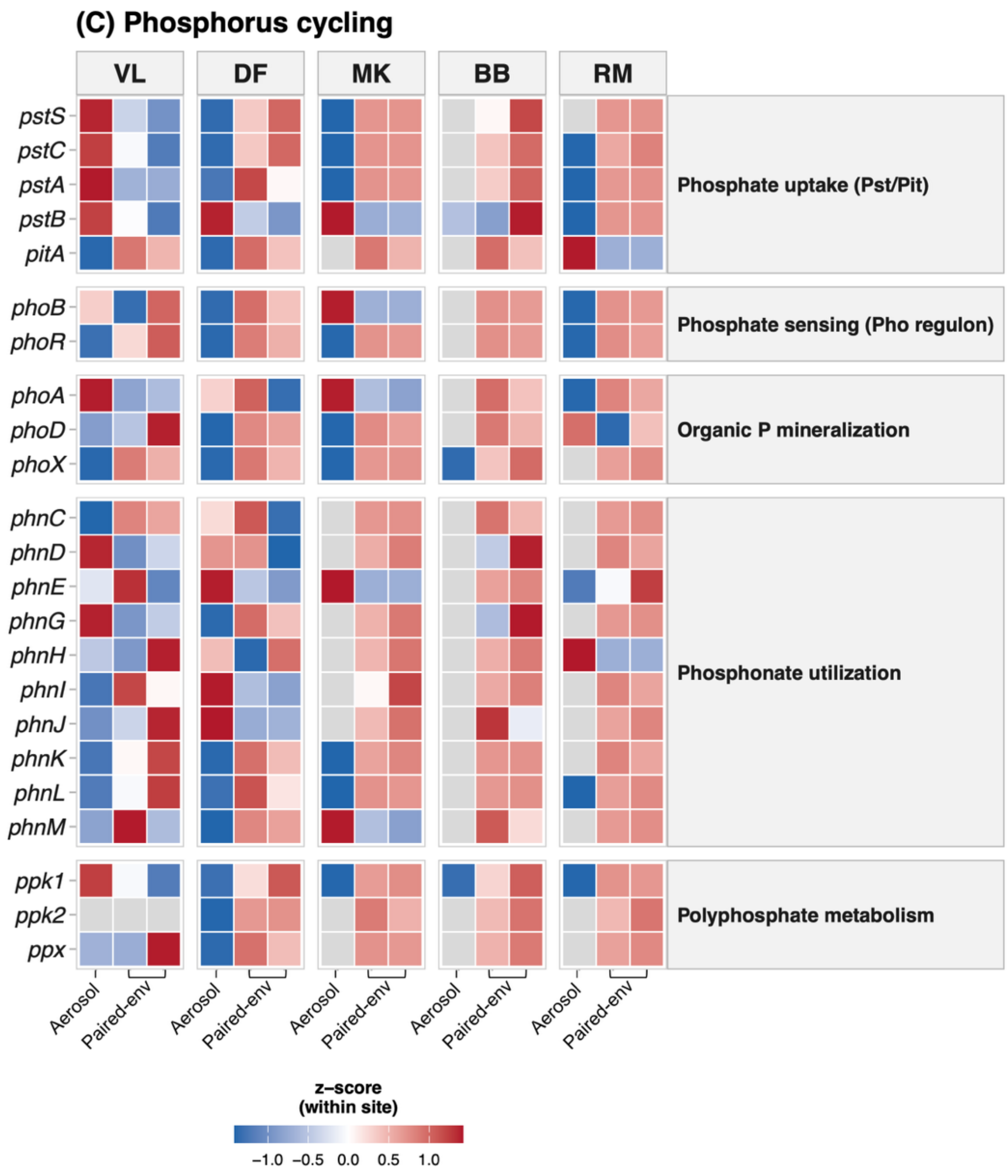


**Supplementary Figure 8 | Phosphorus cycling gene potential across aerosol and paired environmental samples.** As Supplementary Figure 5.

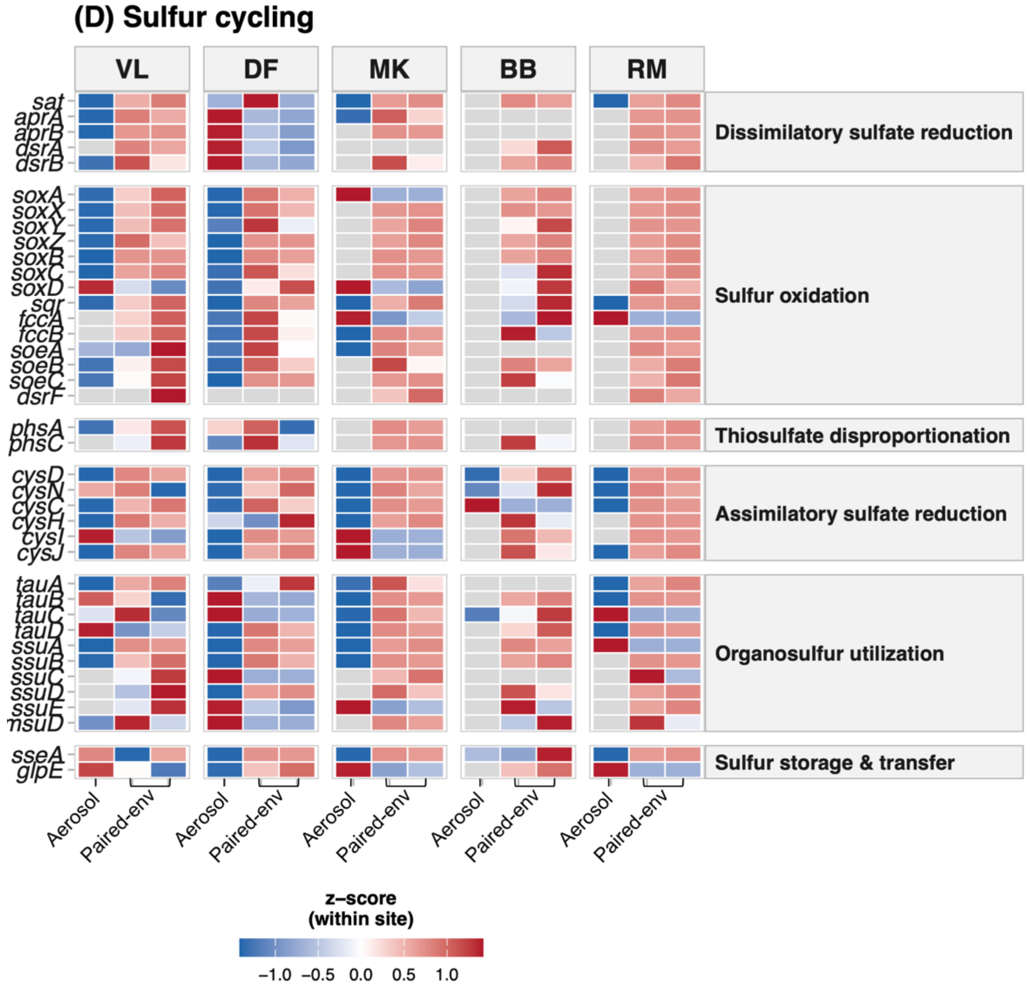


**Supplementary Figure 9 | Sulfur cycling gene potential across aerosol and paired environmental samples.** As Supplementary Figure 5.

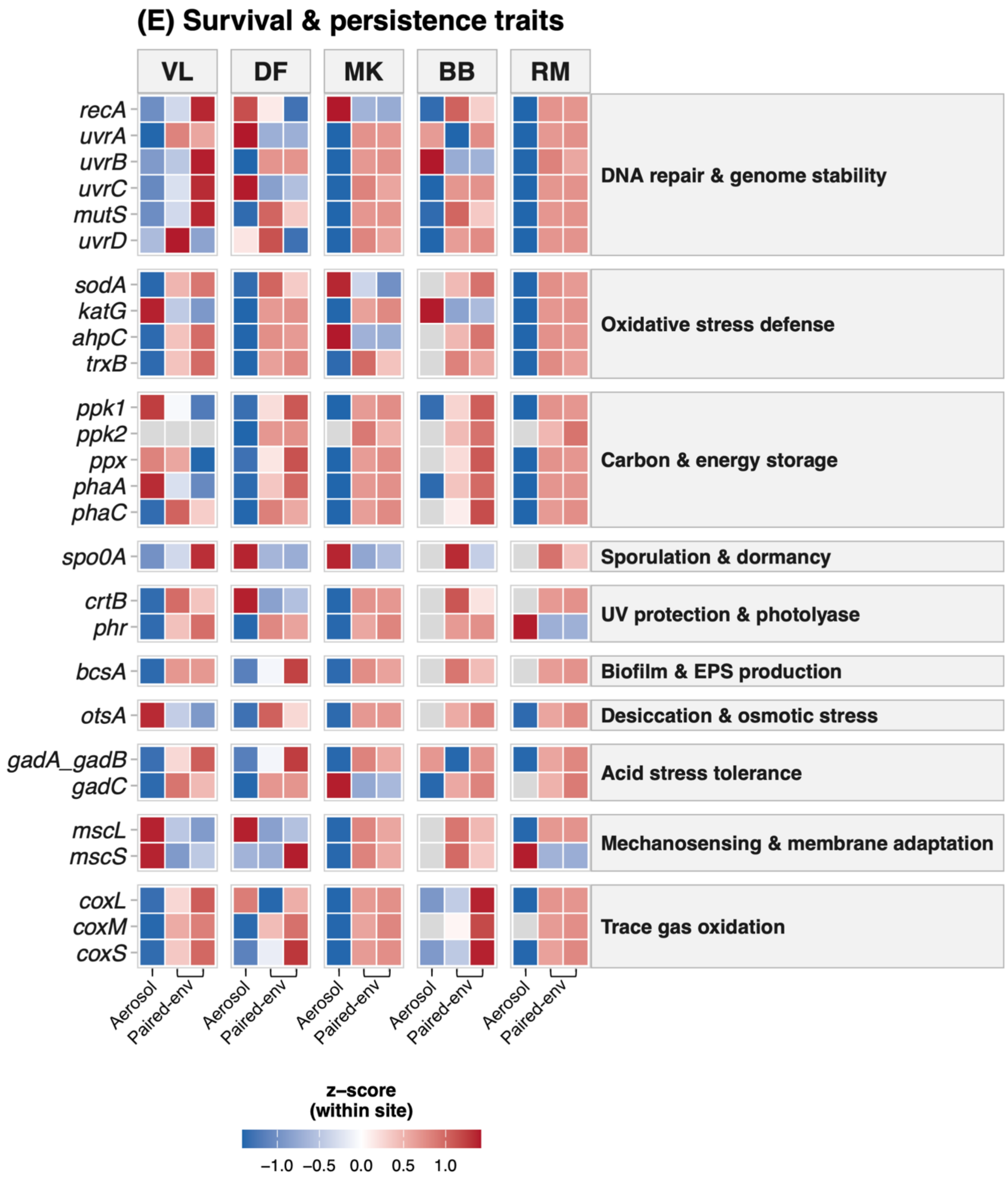


**Supplementary Figure 10 | Survival and persistence trait gene potential across aerosol and paired environmental samples.** As Supplementary Figure 5.

**Supplementary Table S1.** Curated marker gene framework used for gene-centric functional profiling. Marker genes are organized by functional module and mapped to KEGG Orthology (KO) identifiers. All genes were selected a priori based on canonical biochemical pathways independently of their detection in the dataset. Categories span C1 and trace gas metabolism, complex VOC and multi-carbon degradation, nitrogen cycling, phosphorus cycling, sulfur cycling, and survival- and persistence-related traits.

### Nitrogen cycling: Functional categories and canonical genes

| **Category** | **Process** | **Key marker genes (KO)** |
|---|---|---|
| Nitrogen fixation | $N_2 \rightarrow NH_3$ | nifH (K02588), nifD (K02586), nifK (K02591) |
| Ammonia oxidation | $NH_3 \rightarrow NO_2^-$ | amoA (K10944), amoB (K10945), amoC (K10946); hao (K10535) |
| Nitrite oxidation | $NO_2^- \rightarrow NO_3^-$ | nxrA (K00370), nxrB (K00371) |
| Denitrification | $NO_3^- \rightarrow N_2$ | narI (K00374); napA (K02567), napB (K02568); nirK (K00368) / nirS (K15864); norB (K04561), norC (K02305); nosZ (K00376) |
| DNRA | $NO_3^- \rightarrow NH_4^+$ | nrfA (K03385), nrfH (K15876) |
| Assimilatory nitrate reduction | Uptake for biomass | nasA (K00372), nirA (K00366) |
| Anaerobic ammonium oxidation | $NH_4^+ + NO_2^- \rightarrow N_2$ | hzsA (K20932), hzsB (K20933), hzsC (K20934); hzo (K20935) |
| Urea hydrolysis & transport | Urea $\rightarrow NH_3$ | ureA (K01428), ureB (K01429), ureC (K01430); urtA (K11959), urtB (K11960), urtC (K11961), urtD (K11962), urtE (K11963) |

### Phosphate cycling: Functional categories and canonical genes

| **Category** | **Process** | **Marker genes (KO)** |
|---|---|---|
| Phosphate uptake (Pst/Pit) | Inorganic phosphate acquisition | pstS (K02040), pstC (K02038), pstA (K02037), pstB (K02036); pitA (K01083) |

| | | |
|---|---|---|
| Phosphate sensing (Pho regulon) | P starvation regulation | phoB (K07657), phoR (K07636) |
| Organic P mineralization | Release Pi from organic P | phoA (K01077), phoD (K01113), phoX (K01114) |
| Phosphonate utilization | Use of C–P compounds | phnC (K02044), phnD (K02043), phnE (K02042); phnG (K06163), phnH (K06164), phnI (K06165), phnJ (K06166), phnK (K06167), phnL (K06168), phnM (K06169) |
| Polyphosphate metabolism | P storage and buffering | ppk1 (K00937), ppk2 (K01093), ppx (K01095) |

### C1 metabolism

| **Category** | **Process** | **Key marker genes (KO)** |
|---|---|---|
| Methane oxidation | $CH_4 \rightarrow CH_3OH$ | pmoC (K10946), pmoA (K10944), pmoB (K10945); mmoX (K16157), mmoY (K16158), mmoZ (K16159), mmoD (K16162), mmoC (K16161) |
| Methanol oxidation | $CH_3OH \rightarrow HCHO$ | xoxF (K10823), mxaF (K14028), mxaI (K14029) |
| Formaldehyde processing | Detoxification + branch metabolism | fae (K10798), fch (K01455), mch (K08691), folD (K01491), frmA (K00122), frmB (K01468) |
| Formate oxidation | $HCOO^- \rightarrow CO_2$ | fdhA (K00123), fdhB (K00124), fdhC (K00125) |
| C1 assimilation (RuMP) | Formaldehyde fixation | hps (K13812), phi (K02331) |
| C1 assimilation (Serine cycle) | Formaldehyde fixation | glyA (K00600), mcl (K01638), sga (K01783), mtkA (K00614), mtkB (K00615) |
| CO oxidation | $CO \rightarrow CO_2$ | coxL (K03518), coxM (K03519), coxS (K03520) |
| Calvin cycle | Autotrophic carbon fixation | rbcL (K01601), rbcS (K01602), prkB (K00855) |

| | | |
|---|---|---|
| Hydrogen metabolism | | hoxK (K06282), hoxG (K06281), hoxZ (K06283), hupS (K18016), hupL (K18017), hupC (K18018), hoxF (K06284), hoxU (K06285), hoxY (K06286), hoxH (K06287), hoxB (K06288), hoxC (K06289), hoxJ (K06290), hoxA (K06291), hhyS (K22622), hhyL (K22623), hypA (K04651), hypB (K04652), hypC (K04653), hypD (K04654), hypE (K04655), hypF (K04656), hupD (K06280), hydA (K00533) |

### Complex VOC / multi-carbon metabolism

| **Category** | **Process** | **Key marker genes (KO)** |
|---|---|---|
| Alcohol & aldehyde oxidation | Short-chain VOC turnover | adh (K13953), aldh (K00128) |
| Organic acid metabolism | Acetate, lactate, pyruvate use | acs (K01895), ldh (K00016), poxB (K00156) |
| Aromatic compound degradation | Benzoate, catechol, lignin derivatives | benA (K04110), benB (K04111), benC (K04112); catA (K03381), catB (K03382), catC (K03383); pcaG (K00448), pcaH (K00449) |
| Alkane & hydrocarbon degradation | Alkane / alkene oxidation | alkB (K00496), ladA (K16244), cyp153 (K15446) |
| Terpene degradation | Biogenic VOCs | atuA (K06443), atuB (K06444), limA (K07539) |
| Carbohydrate degradation | Polymer breakdown | celA (K01179), xynA (K01181), amyA (K01176) |

### Sulfur cycling: Functional categories and marker genes

| **Category** | **Process** | **Key marker genes (KO)** |
|---|---|---|
| Dissimilatory sulfate reduction | $SO_4^{2-} \rightarrow H_2S$ | sat (K00958), aprA (K00394), aprB (K00395), dsrA (K11180), dsrB (K11181) |

| | | |
|---|---|---|
| Sulfur oxidation | Reduced sulfur → $SO_4^{2-}$ | soxA (K17222), soxX (K17223), soxY (K17224), soxZ (K17225), soxB (K17226), soxC (K17227), soxD (K17228); sqr (K17218); fccA (K15228), fccB (K15229); soeA (K21307), soeB (K21308), soeC (K21309), dsrC (K17217), dsrE (K17219), dsrF (K17220), dsrH (K17221) |
| Thiosulfate disproportionation | $S_2O_3^{2-}$ → $S^0$ / $H_2S$ / $SO_4^{2-}$ | phsA (K08352), phsB (K08353), phsC (K08354) |
| Assimilatory sulfate reduction | $SO_4^{2-}$ → cysteine | cysD (K00955), cysN (K00956), cysC (K00957), cysH (K00392), cysI (K00380), cysJ (K00381) |
| Organosulfur utilization | Sulfonates, organosulfur compounds | tauA (K07239), tauB (K07240), tauC (K07241), tauD (K03118); ssuA (K12339), ssuB (K12340), ssuC (K12341), ssuD (K12342), ssuE (K12343); msuD (K18275); dmdA (K14468) |
| Sulfur storage & transfer | Sulfur trafficking and detoxification | sseA (K00658), glpE (K01011) |

### Survival- and persistence-related functional traits assessed in airborne metagenomes

| **Category** | **Process** | **Key marker genes (KO)** |
|---|---|---|
| DNA repair & genome stability | Repair of UV- and oxidative-induced DNA damage; maintenance of genome integrity | recA (K03553); uvrA (K03701), uvrB (K03702), uvrC (K03703); mutS (K02335); uvrD (K03555) |

| | | |
|---|---|---|
| Oxidative-stress defense | Detoxification of reactive oxygen species and redox homeostasis | sodA (K04564); katG (K03781); ahpC (K03386); trxB (K03671) |
| Carbon & energy storage | Carbon, energy, and phosphate storage during nutrient limitation | ppk1 (K00937); ppk2 (K01093); ppx (K01524); phaA (K00626), phaB (K00025), phaC (K03821) |
| Sporulation & dormancy | Entry into dormant states enabling long-term persistence | spo0A (K07699) |
| UV protection & photolyase | Photoprotection and UV shielding | crtB (K02291); phr (photolyase) (K01669) |
| Biofilm & EPS production | Extracellular matrix production, aggregation, and surface attachment | bcsA (K00694) |
| Desiccation & osmotic stress | Protection against dehydration and osmotic stress via compatible solutes | otsA (K00697) |
| Acid stress tolerance | Cytoplasmic pH homeostasis under acidic conditions | gadA/gadB (K01580); gadC (K01581) |
| Mechanosensing & membrane adaptation | Sensing and relief of membrane tension during osmotic shock | mscL (K03823); mscS (K03824) |
| C1 & trace-gas metabolism | Oxidation of methane, CO, and other trace gases | pmoA (K10944), pmoB (K10945), pmoC (K10946); coxL (K03518), coxM (K03519), coxS (K03520) |

**Supplementary Table S2.** Proportion of classified reads (%) assigned to Bacteria, Archaea, Eukaryota, and Viruses in each sample, based on Kraken2/Bracken classification after removal of human reads.

| Sample | Type | Archaea | Bacteria | Eukaryota | Viruses |
|---|---|---|---|---|---|
| A_BB1 | Aerosol | 0.425 | 81.157 | 18.211 | 0.208 |
| A_DF1 | Aerosol | 0.597 | 98.343 | 1.024 | 0.036 |
| A_MK1 | Aerosol | 0.286 | 20.112 | 79.493 | 0.108 |
| A_RM1 | Aerosol | 0.79 | 69.375 | 29.05 | 0.785 |
| A_VL2 | Aerosol | 0.114 | 97.702 | 2.177 | 0.007 |
| S_DF1 | Paired environment | 0.321 | 99.548 | 0.101 | 0.03 |
| S_DF2 | Paired environment | 0.296 | 99.578 | 0.098 | 0.029 |
| S_MK1 | Paired environment | 0.544 | 98.908 | 0.383 | 0.165 |
| S_MK2 | Paired environment | 0.537 | 98.97 | 0.387 | 0.106 |
| S_RM1 | Paired environment | 0.838 | 98.742 | 0.379 | 0.041 |
| S_RM2 | Paired environment | 0.769 | 98.81 | 0.365 | 0.055 |
| S_VL1 | Paired environment | 0.766 | 98.513 | 0.679 | 0.041 |
| S_VL2 | Paired environment | 0.314 | 99.418 | 0.214 | 0.053 |
| W_BB1 | Paired environment | 0.473 | 98.089 | 1.225 | 0.214 |
| W_BB2 | Paired environment | 0.446 | 96.9 | 2.511 | 0.143 |

**Supplementary Table S3. Genera differentially abundant between aerosol and paired environmental samples.** Bacterial and eukaryotic genera with Benjamini–Hochberg adjusted $p < 0.05$ and $|\log_2$ fold change$| \geq 2$ in site-blocked DESeq2 analysis (Fig. 4). Positive $\log_2$ fold changes (shrunken with apeglm) indicate enrichment in aerosols. baseMean is the mean normalized count across all samples.

| Domain | Genus | Direction | log2FC | padj | baseMean |
|---|---|---|---|---|---|
| Bacteria | Shigella | Depleted in aerosols | -6.75 | 6.87E-10 | 2104.4 |
| Bacteria | Seohaeicola | Depleted in aerosols | -5.94 | 2.03E-09 | 4657.2 |
| Bacteria | Acidithiobacillus | Depleted in aerosols | -4.56 | 3.83E-06 | 503.8 |
| Bacteria | Opitutus | Depleted in aerosols | -2.19 | 4.86E-05 | 15238.8 |
| Bacteria | Horticoccus | Depleted in aerosols | -2.05 | 1.98E-04 | 3236.9 |
| Bacteria | Desulfomicrobium | Depleted in aerosols | -2.47 | 3.21E-04 | 3197.7 |
| Bacteria | Pelobacter | Depleted in aerosols | -2.14 | 0.00147 | 6113.4 |
| Bacteria | Staphylococcus | Enriched in aerosols | 5.77 | 2.00E-76 | 306165.3 |
| Bacteria | Sellimonas | Enriched in aerosols | 4.3 | 6.76E-75 | 4366.4 |
| Bacteria | Mammaliicoccus | Enriched in aerosols | 6.8 | 7.34E-68 | 33931.6 |
| Bacteria | Jeotgalicoccus | Enriched in aerosols | 5.02 | 3.75E-53 | 25101 |
| Bacteria | Ligilactobacillus | Enriched in aerosols | 5.17 | 5.26E-39 | 7148.8 |
| Bacteria | Weissella | Enriched in aerosols | 4.2 | 1.55E-31 | 6486.7 |
| Bacteria | Romboutsia | Enriched in aerosols | 4.18 | 3.89E-26 | 23748.7 |
| Bacteria | Caldifermentibacillus | Enriched in aerosols | 2.99 | 6.86E-24 | 4460 |
| Bacteria | Claveliimonas | Enriched in aerosols | 2.8 | 1.37E-21 | 1320.8 |
| Bacteria | Lactobacillus | Enriched in aerosols | 4.62 | 3.07E-21 | 23756.9 |
| Bacteria | Nosocomiicoccus | Enriched in aerosols | 5.22 | 3.58E-19 | 7443.2 |
| Bacteria | Subdoligranulum | Enriched in aerosols | 2.44 | 3.75E-18 | 4195.8 |

| | | | | | |
|---|---|---|---|---|---|
| Bacteria | Aerococcus | Enriched in aerosols | 5.65 | 1.25E-17 | 226464.3 |
| Bacteria | Leuconostoc | Enriched in aerosols | 3.78 | 2.07E-14 | 10169.1 |
| Bacteria | Ruoffia | Enriched in aerosols | 4.61 | 3.15E-14 | 13112.9 |
| Bacteria | Limosilactobacillus | Enriched in aerosols | 2.98 | 4.84E-14 | 5882.7 |
| Bacteria | Lacticaseibacillus | Enriched in aerosols | 2.95 | 2.75E-13 | 3508.8 |
| Bacteria | Wolbachia | Enriched in aerosols | 5.35 | 1.20E-12 | 11081.1 |
| Bacteria | Ureibacillus | Enriched in aerosols | 3.54 | 1.25E-12 | 2837.3 |
| Bacteria | Levilactobacillus | Enriched in aerosols | 3.17 | 2.95E-12 | 1137.7 |
| Bacteria | Carnobacterium | Enriched in aerosols | 4.21 | 3.02E-12 | 84723.3 |
| Bacteria | Brevibacterium | Enriched in aerosols | 2.95 | 6.15E-12 | 78880.8 |
| Bacteria | Brachybacterium | Enriched in aerosols | 3.81 | 3.78E-11 | 137457.5 |
| Bacteria | Fructilactobacillus | Enriched in aerosols | 4.04 | 3.78E-11 | 712 |
| Bacteria | Streptococcus | Enriched in aerosols | 2.7 | 6.56E-11 | 38954.6 |
| Bacteria | Brochothrix | Enriched in aerosols | 3.61 | 1.53E-10 | 1352.9 |
| Bacteria | Peribacillus | Enriched in aerosols | 3.79 | 3.56E-10 | 20789.1 |
| Bacteria | Lactococcus | Enriched in aerosols | 2.66 | 3.76E-10 | 45380.6 |
| Bacteria | Turicibacter | Enriched in aerosols | 3.22 | 1.15E-09 | 6866.4 |
| Bacteria | Priestia | Enriched in aerosols | 3 | 2.07E-09 | 11279.2 |
| Bacteria | Massilistercora | Enriched in aerosols | 2.03 | 5.66E-09 | 1724.6 |
| Bacteria | Tetragenococcus | Enriched in aerosols | 2.36 | 6.85E-09 | 4851.9 |
| Bacteria | Megamonas | Enriched in aerosols | 3.16 | 7.98E-09 | 1010.4 |
| Bacteria | Paraclostridium | Enriched in aerosols | 3.47 | 1.79E-08 | 10231.3 |
| Bacteria | Macrococcoides | Enriched in aerosols | 3.56 | 1.86E-08 | 1458.8 |
| Bacteria | Psychrobacter | Enriched in aerosols | 3.14 | 4.42E-08 | 59109.8 |
| Bacteria | Bifidobacterium | Enriched in aerosols | 2.27 | 4.77E-08 | 62887.8 |
| Bacteria | Vagococcus | Enriched in aerosols | 2.65 | 8.49E-08 | 3595.2 |
| Bacteria | Frigoribacterium | Enriched in aerosols | 2.26 | 1.71E-07 | 13115.4 |

| | | | | | |
|---|---|---|---|---|---|
| Bacteria | Thermoactinomyces | Enriched in aerosols | 2.61 | 2.25E-07 | 1583.5 |
| Bacteria | Blautia | Enriched in aerosols | 2.73 | 2.44E-07 | 16708.4 |
| Bacteria | Terrisporobacter | Enriched in aerosols | 3.43 | 2.47E-07 | 5146.6 |
| Bacteria | Heyndrickxia | Enriched in aerosols | 2.51 | 4.65E-07 | 2380.6 |
| Bacteria | Intestinibacter | Enriched in aerosols | 3.29 | 5.11E-07 | 1034.4 |
| Bacteria | Sarcina | Enriched in aerosols | 3.18 | 7.17E-07 | 837.3 |
| Bacteria | Planococcus | Enriched in aerosols | 3.06 | 7.83E-07 | 9066 |
| Bacteria | Lactiplantibacillus | Enriched in aerosols | 3.67 | 8.52E-07 | 23628.7 |
| Bacteria | Marinilactibacillus | Enriched in aerosols | 3.32 | 8.52E-07 | 2418.5 |
| Bacteria | Neomicrococcus | Enriched in aerosols | 3.76 | 1.39E-06 | 96534.4 |
| Bacteria | Ruminococcus | Enriched in aerosols | 2.35 | 1.41E-06 | 68365.2 |
| Bacteria | Metamycoplasma | Enriched in aerosols | 3.63 | 1.80E-06 | 2728.8 |
| Bacteria | Cetobacterium | Enriched in aerosols | 3.34 | 1.87E-06 | 1410.6 |
| Bacteria | Erwinia | Enriched in aerosols | 2.67 | 1.93E-06 | 14199.8 |
| Bacteria | Ruminiclostridium | Enriched in aerosols | 2.02 | 2.28E-06 | 981.8 |
| Bacteria | Helcococcus | Enriched in aerosols | 3.41 | 3.13E-06 | 3008.7 |
| Bacteria | Allocoprobacillus | Enriched in aerosols | 2.97 | 3.32E-06 | 826.5 |
| Bacteria | Enterococcus | Enriched in aerosols | 2.74 | 3.32E-06 | 177207.3 |
| Bacteria | Corynebacterium | Enriched in aerosols | 2.52 | 3.56E-06 | 1154096.8 |
| Bacteria | Friedmanniella | Enriched in aerosols | 2.16 | 3.69E-06 | 7805.2 |
| Bacteria | Stanieria | Enriched in aerosols | 3.3 | 4.69E-06 | 2077.1 |
| Bacteria | Companilactobacillus | Enriched in aerosols | 3.19 | 6.25E-06 | 614.7 |
| Bacteria | Fusobacterium | Enriched in aerosols | 3.13 | 6.25E-06 | 14662.7 |
| Bacteria | Virgibacillus | Enriched in aerosols | 3.08 | 8.06E-06 | 2471.5 |
| Bacteria | Veillonella | Enriched in aerosols | 3.14 | 1.09E-05 | 1084.2 |
| Bacteria | Paenibacillus | Enriched in aerosols | 2.38 | 1.45E-05 | 12310.2 |
| Bacteria | Thermobacillus | Enriched in aerosols | 2.67 | 1.45E-05 | 7459.7 |

| | | | | | |
|---|---|---|---|---|---|
| Bacteria | Thomasclavelia | Enriched in aerosols | 2.51 | 1.85E-05 | 1844.7 |
| Bacteria | Gottschalkia | Enriched in aerosols | 3.22 | 1.96E-05 | 2282.6 |
| Bacteria | Lacrimispora | Enriched in aerosols | 2.55 | 1.98E-05 | 1874 |
| Bacteria | Anaerostipes | Enriched in aerosols | 2.33 | 2.19E-05 | 2308.4 |
| Bacteria | Micrococcus | Enriched in aerosols | 2.34 | 2.19E-05 | 72463 |
| Bacteria | Vallitalea | Enriched in aerosols | 2.85 | 2.61E-05 | 1618.7 |
| Bacteria | Metabacillus | Enriched in aerosols | 2.8 | 2.94E-05 | 964.3 |
| Bacteria | Plantibacter | Enriched in aerosols | 2.07 | 2.94E-05 | 11387.2 |
| Bacteria | Oceanirhabdus | Enriched in aerosols | 2.78 | 3.19E-05 | 1378.8 |
| Bacteria | Tepidibacter | Enriched in aerosols | 2.53 | 3.19E-05 | 1408 |
| Bacteria | Clostridium | Enriched in aerosols | 2.25 | 3.83E-05 | 97600.3 |
| Bacteria | Brachyspira | Enriched in aerosols | 2.85 | 3.85E-05 | 6151.8 |
| Bacteria | Desemzia | Enriched in aerosols | 2.93 | 3.85E-05 | 9715.9 |
| Bacteria | Agathobacter | Enriched in aerosols | 2.6 | 4.07E-05 | 2339.8 |
| Bacteria | Ehrlichia | Enriched in aerosols | 2.88 | 4.27E-05 | 1957.9 |
| Bacteria | Halotia | Enriched in aerosols | 2.03 | 4.53E-05 | 926.4 |
| Bacteria | Gloeothece | Enriched in aerosols | 2.82 | 5.06E-05 | 1182.1 |
| Bacteria | Hathewaya | Enriched in aerosols | 2.7 | 5.15E-05 | 671.1 |
| Bacteria | Globicatella | Enriched in aerosols | 2.31 | 5.34E-05 | 1079.6 |
| Bacteria | Fervidibacillus | Enriched in aerosols | 2.93 | 7.96E-05 | 1851.2 |
| Bacteria | Crinalium | Enriched in aerosols | 2.32 | 8.58E-05 | 629.7 |
| Bacteria | Anaerocolumna | Enriched in aerosols | 2.02 | 8.76E-05 | 1845.4 |
| Bacteria | Crassaminicella | Enriched in aerosols | 2.37 | 8.76E-05 | 695.4 |
| Bacteria | Lentilactobacillus | Enriched in aerosols | 2.36 | 9.57E-05 | 1907.3 |
| Bacteria | Prochlorococcus | Enriched in aerosols | 2.7 | 1.08E-04 | 19281.3 |
| Bacteria | Mesomycoplasma | Enriched in aerosols | 2.74 | 1.38E-04 | 2710.3 |
| Bacteria | Gracilibacillus | Enriched in aerosols | 2.44 | 1.49E-04 | 579.8 |

| | | | | | |
|---|---|---|---|---|---|
| Bacteria | Psychrobacillus | Enriched in aerosols | 2.64 | 1.53E-04 | 1501.9 |
| Bacteria | Brevibacillus | Enriched in aerosols | 2.31 | 1.63E-04 | 992 |
| Bacteria | Chryseobacterium | Enriched in aerosols | 2.19 | 1.70E-04 | 53200.9 |
| Bacteria | Lysinibacillus | Enriched in aerosols | 2.23 | 1.73E-04 | 5637.8 |
| Bacteria | Mycoplasma | Enriched in aerosols | 2.72 | 1.94E-04 | 4935.7 |
| Bacteria | Tepiditoga | Enriched in aerosols | 2.54 | 1.97E-04 | 539.8 |
| Bacteria | Alkaliphilus | Enriched in aerosols | 2.3 | 2.06E-04 | 668.9 |
| Bacteria | Gemella | Enriched in aerosols | 2.39 | 2.06E-04 | 591.4 |
| Bacteria | Chroococcidiopsis | Enriched in aerosols | 2.3 | 2.18E-04 | 3439.7 |
| Bacteria | Finegoldia | Enriched in aerosols | 2.51 | 2.18E-04 | 1509.9 |
| Bacteria | Thermoanaerobacterium | Enriched in aerosols | 2.36 | 2.18E-04 | 2879.6 |
| Bacteria | Acetobacteroides | Enriched in aerosols | 2.41 | 2.18E-04 | 9297.2 |
| Bacteria | Halanaerobium | Enriched in aerosols | 2.46 | 2.28E-04 | 1601.5 |
| Bacteria | Spiroplasma | Enriched in aerosols | 2.75 | 3.15E-04 | 8677 |
| Bacteria | Caloramator | Enriched in aerosols | 2.54 | 3.21E-04 | 5033.2 |
| Bacteria | Trichodesmium | Enriched in aerosols | 2.52 | 3.31E-04 | 4043.5 |
| Bacteria | Haloimpatiens | Enriched in aerosols | 2.19 | 3.35E-04 | 1627.9 |
| Bacteria | Candidatus Arthromitus | Enriched in aerosols | 2.83 | 4.05E-04 | 703 |
| Bacteria | Faecalibacillus | Enriched in aerosols | 2.39 | 4.31E-04 | 1606.7 |
| Bacteria | Peptacetobacter | Enriched in aerosols | 2.32 | 4.55E-04 | 1251.3 |
| Bacteria | Oceanobacillus | Enriched in aerosols | 2.14 | 4.88E-04 | 2309.4 |
| Bacteria | Parvimonas | Enriched in aerosols | 2.1 | 5.20E-04 | 1808 |
| Bacteria | Campylobacter | Enriched in aerosols | 2.21 | 5.50E-04 | 13518.7 |
| Bacteria | Ilyobacter | Enriched in aerosols | 2.05 | 5.73E-04 | 911.2 |
| Bacteria | Butyrivibrio | Enriched in aerosols | 2.08 | 5.83E-04 | 5455.5 |
| Bacteria | Latilactobacillus | Enriched in aerosols | 2.14 | 6.21E-04 | 3505.4 |
| Bacteria | Anaeromicropila | Enriched in aerosols | 2.52 | 6.30E-04 | 2563.4 |

| | | | | | |
|---|---|---|---|---|---|
| Bacteria | Marnyiella | Enriched in aerosols | 2.2 | 6.30E-04 | 1123.6 |
| Bacteria | Tissierella | Enriched in aerosols | 2.11 | 7.05E-04 | 771.6 |
| Bacteria | Mycoplasmopsis | Enriched in aerosols | 2.45 | 7.44E-04 | 3856 |
| Bacteria | Thermobaculum | Enriched in aerosols | 2.44 | 7.77E-04 | 4726.3 |
| Bacteria | Planktothrix | Enriched in aerosols | 2.06 | 8.20E-04 | 1032 |
| Bacteria | Gloeocapsopsis | Enriched in aerosols | 2.42 | 8.24E-04 | 7659.3 |
| Bacteria | Novibacillus | Enriched in aerosols | 2.06 | 0.00103 | 1472.1 |
| Bacteria | Candidatus Hodgkinia | Enriched in aerosols | 2.33 | 0.0011 | 1682 |
| Bacteria | Trichormus | Enriched in aerosols | 2.27 | 0.0011 | 1156.1 |
| Bacteria | Orientia | Enriched in aerosols | 2.09 | 0.00114 | 763.4 |
| Bacteria | Oscillatoria | Enriched in aerosols | 2.06 | 0.00125 | 5309.1 |
| Bacteria | Proteinivorax | Enriched in aerosols | 2.21 | 0.00149 | 962.3 |
| Bacteria | Miniphocaeibacter | Enriched in aerosols | 2.39 | 0.00156 | 627.2 |
| Bacteria | Jeotgalibaca | Enriched in aerosols | 2.21 | 0.00169 | 36283.2 |
| Bacteria | Kaistella | Enriched in aerosols | 2.08 | 0.00179 | 21570.7 |
| Bacteria | Amedibacillus | Enriched in aerosols | 2.01 | 0.00184 | 1374.6 |
| Bacteria | Okeanomitos | Enriched in aerosols | 2.14 | 0.00191 | 7489.7 |
| Bacteria | Macrococcus | Enriched in aerosols | 2.04 | 0.00259 | 1328.8 |
| Bacteria | Caminicella | Enriched in aerosols | 2.07 | 0.00396 | 948.5 |
| Bacteria | Geobacillus | Enriched in aerosols | 2.03 | 0.00426 | 4399.7 |
| Bacteria | Anaerobutyricum | Enriched in aerosols | 2.13 | 0.00432 | 2115.9 |
| Eukaryota | Leishmania | Depleted in aerosols | -2.17 | 3.29E-16 | 7228 |
| Eukaryota | Besnoitia | Depleted in aerosols | -2.13 | 6.79E-10 | 1665.1 |
| Eukaryota | Marasmius | Enriched in aerosols | 5.59 | 1.53E-145 | 17986 |
| Eukaryota | Psilocybe | Enriched in aerosols | 6.23 | 3.95E-137 | 39130.5 |
| Eukaryota | Rhizoctonia | Enriched in aerosols | 5.15 | 5.25E-95 | 12450 |
| Eukaryota | Botrytis | Enriched in aerosols | 3.5 | 6.79E-10 | 26112.5 |

| | | | | | |
|---|---|---|---|---|---|
| Eukaryota | Debaryomyces | Enriched in aerosols | 2.44 | 9.76E-10 | 3754.2 |
